# Adaptations of DNA viruses are influenced by host and environment with proliferations constrained by environmental niche

**DOI:** 10.1101/2025.07.28.667335

**Authors:** Michael Hoggard, Emilie Gios, Hwee Sze Tee, Jemma L. Geoghegan, Kim M. Handley

## Abstract

Viruses are ubiquitous albeit individually constrained by host-range. Less well understood are environmental limitations on virus proliferation. To investigate estuarine viral diversity, niche constraints, and genomic traits of environmental adaptation, we analysed metagenomic and metatranscriptomic data from across an estuarine salinity gradient, including water and sediment habitats. We then expanded our analysis to globally distributed viral genomes. Viral distributions varied by estuary habitat, reflecting prokaryote community patterns, and highlighting that virus-host interactions are strongly influenced by environment. Viral lineages, up until approximately the rank of genus, were largely partitioned by ecological niche based on factors such as salinity and the aquatic-terrestrial divide. Across habitat boundaries, we identified variations in genomic GC content, and genomic signatures of salinity adaptation, particularly in virus major tail and capsid proteins. These included elevated ratios of acidic to basic amino acids and decreased protein isoelectric points at higher salinities which did not solely reflect biases inherited due to reliance on host machinery. Salinity is a well-described physiological stressor for prokaryotes. Results suggest that viruses employ some of the same osmoadaptive strategies independent of adaptation based on host machinery. Our findings indicate that, in addition to host range limitations, successful proliferation of viruses into distinct biomes (e.g. freshwater, saline, terrestrial) is constrained by adaptation to specific ecological niches.

## INTRODUCTION

Viruses are globally ubiquitous and abundant^1,2^ and exert a strong effect on microbial communities. This includes influencing microbial evolution^3^, community diversity^4^ and turnover^5^, metabolic reprogramming of infected cells^6–10^, horizontal gene transfer^4^, and as an important modifier of nutrient flux and biogeochemical cycling in ecosystems globally^4,9,11,12^. Virus-mediated microbial lysis can be considerable, accounting for up to 27% of prokaryote cell death in the open ocean^5^. In addition to shaping microbial community structure and diversity, the release of dissolved organic matter during cell lysis reduces nutrient flow to higher trophic levels (via grazing of microbes), modifies sequestration into the deep sea (due to particle aggregation and sinking via the biological pump and viral shuttle)^4,9^, and recycles nutrients back into local microbial growth, metabolism, and primary production (viral shunt)^4,5,9,10,12–14^. These virus-mediated processes affect global primary productivity and nutrient cycling at scales relevant to estimations of global carbon sequestration^5,12^.

Differences in environmental factors, such as salinity, pH and temperature, or substrate type (e.g., sediment, water, or host), are important variables governing the distribution and diversity of prokaryotes^15–18^. Adaptations related to these differences are reflected in genomic traits, for example traits associated with salinity (e.g., encoded amino acid modifications for protein stability and osmoregulatory genes)^15,19,20^ and temperature (e.g., genome-streamlining, GC content, and again, encoded amino acid modifications)^21^. These adaptations can confine prokaryotes to a set niche. For example, niche transitions between freshwater and saline environments are rare and evolutionarily ancient^22,23^. Environmental differentiation of DNA viral populations is also commonly observed, including by temperature, salinity, and/or oxygen concentration^24–26^, broad “ecological zones”^27^, ocean depth^28,29^, and between aquatic and terrestrial habitats^30^. This differentiation has been shown to mirror that of putative prokaryotic host communities^26,29^, and environmental adaptation leading to niche restriction of the putative hosts of viruses is a well-described constraint on the diversity and distribution of viruses in distinct ecosystems^27,31^. Viral adaptation is known to be influenced by host defense mechanisms^32,33^, cell surface receptors^34^, and host translational machinery^35^. However, the extent to which proliferation across habitat boundaries is additionally restricted due to direct adaptation of viruses to the physicochemistry of distinct niches is less well understood. Here, we sought to examine the influence of ecological boundaries and niche partitioning on viral diversification, including whether GC content and amino acid usage biases in viruses are also influenced directly by environmental adaptation.

Estuaries are an interface and mixing zone of distinct aquatic ecosystems, enabling the assessment of viral population distributions, proliferation, and adaptive traits across interconnected habitat boundaries. To investigate the diversity, distribution, and niche restriction of estuarine DNA viruses we analysed metagenomic and metatranscriptomic datasets from across a freshwater-brackish-marine estuarine salinity gradient, including both water and sediment habitats. These ecological attributes of viruses were analysed alongside the distributions of putative hosts, changes in host translational machinery, and genomic traits of viral environmental adaptation. We then expanded our analysis to include all high-quality viral genomes in the IMG/VR database, to assess whether patterns observed in this estuarine dataset were also reflected in a large global database of viruses from diverse ecosystems.

## RESULTS AND DISCUSSION

### *Caudoviricetes* dominate the estuarine habitats

We identified 70,326 putatively viral contigs (each >3,000 bp long) from water and sediment samples (spanning a 0.2 to 35 ppt salinity gradient) (**Table S1**). These were clustered into 31,711 viral operational taxonomic units (vOTUs), with each representing a population of closely related viral sequences. The number of viral contigs clustered per vOTU ranged from 1 to 176 (median=9; interquartile range=4-15), indicating some microdiversity within estuarine vOTUs (**Figure S1**). The subset of vOTUs greater than 50% complete (*n*=858) ranged from 3,079 to 531,516 bp in length (mean 47,351 bp) (**Table S2**). This size distribution is within the range of the estimated full genome sizes of high quality and complete viral genomes in the IMG/VR database (370 to 2,473,870 bp; mean 40,795 bp). Downstream analyses were based on viral contigs or vOTUs >50% complete unless otherwise stated.

In previous metagenomics-based studies, the majority (>90%) of viral sequences have often remained unclassified, generally due to a lack of clustering with known references in the viralRefSeq database. Of those classified, *Caudoviricetes* (formerly *Caudovirales*) are frequently the most common DNA viruses recovered from a broad array of environments, including marine^24,27,36^, groundwater^37^, estuarine^25,26,38^, and soil^30,39,40^. In this study, less than 7% of the final filtered set of 858 vOTUs clustered closely enough with viralRefSeq references to confidently predict genus-level taxonomy (**Table S3**). Final taxonomy predictions for vOTUs were subsequently generated based on similarity to both viralRefSeq and IMG/VR aquatic sequences by, in order, vConTACT2 clustering with viralRefSeq sequences, manual inspection of protein-sharing network interactions, and clustering with high-quality IMG/VR sequences. Fifty-three percent (*n*=452) of estuarine vOTUs clustered together with IMG/VR sequences into 280 separate clusters. Based on the tuning parameters for clustering, sequences within individual clusters are predicted to approximate genus-level relatedness^41^.

Therefore, over half of our vOTUs share a putative genus-level similarity to aquatic viruses in the global IMG/VR database. Network interactions outside of clusters also suggested that most vOTUs shared some relatedness to high-quality viruses in IMG/VR at a rank higher than genus. Furthermore, the combined approach assigned class predictions to 73% of vOTUs (**Table S3**). *Caudoviricetes* (tailed dsDNA phage; phylum *Uroviricota*) were the dominant class overall (*n*=616), followed by *Malgrandaviricetes* (*n*=12, phylum *Phixviricota*) (**Figure 1; Table S2; Table S4**). This distribution is similar to the IMG/VR database, where 72% of high-quality viral sequences are classified as *Caudoviricetes*.

**Figure 1.**
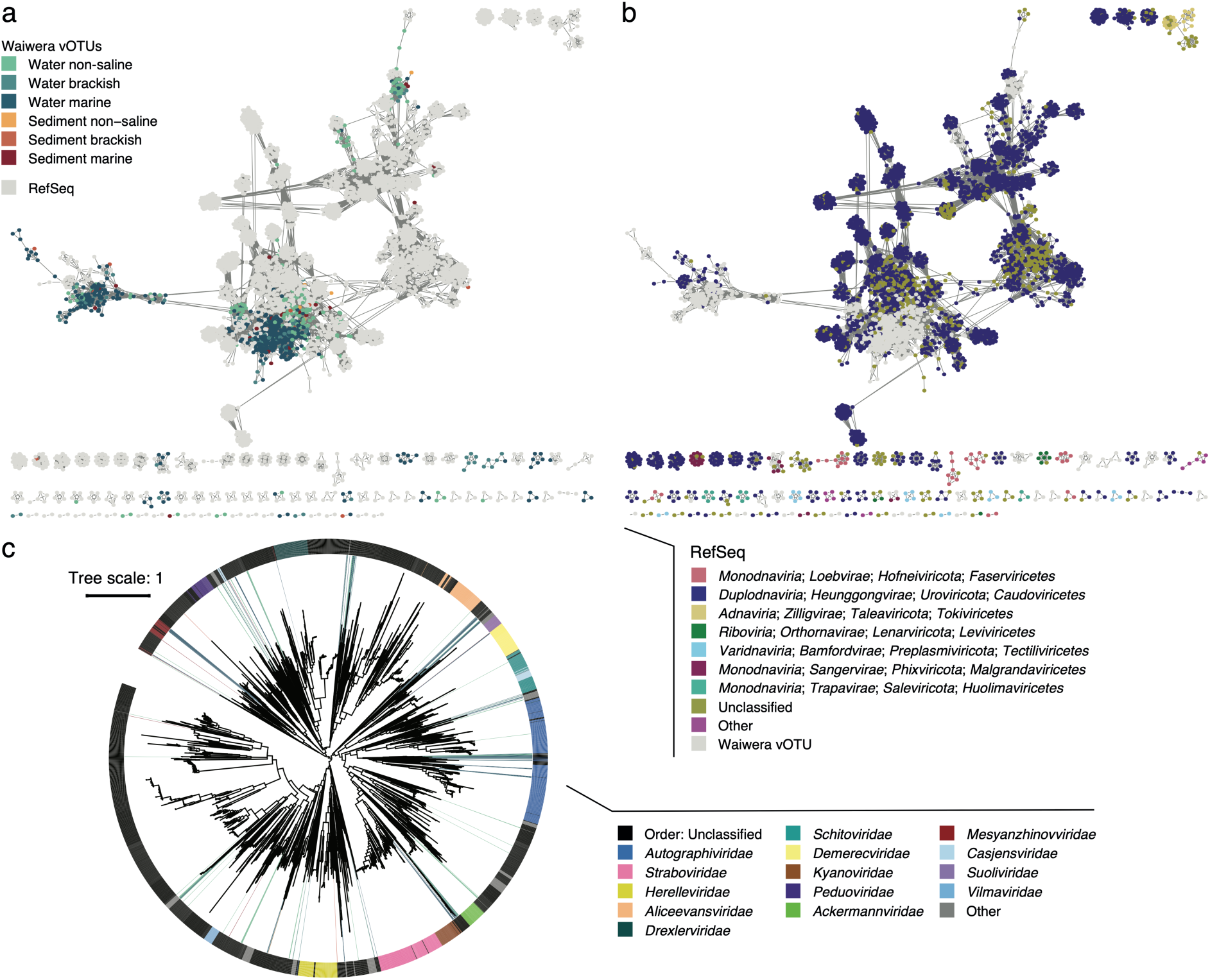
vOTU taxonomy. (a-b) vConTACT2 protein-sharing networks for Waiwera vOTUs and viralRefSeq (v211) coloured by (a) the estuary zone where the vOTU was the most abundant, and (b) viral reference taxonomy. (c) Inferred phylogeny for *Caudoviricetes* vOTUs (205 vOTUs >85% complete) and viralRefSeq references based on concatenated protein alignment of a set of predicted *Caudoviricetes* core genes (*n*=44). Tree generated via IQ-TREE using ModelFinder (with VT+F+I+G4 selected). Lines extending from branch tips for Waiwera vOTUs are coloured by estuary zone (as for part a), and the outer ring is coloured by predicted vOTU and RefSeq taxonomy. The scale bar represents amino acid substitutions per site.

Phylogeny was also inferred for our estuarine *Caudoviricetes* viral genomes (>85% complete; *n*=205) via alignment of concatenated putative single copy core genes. In this analysis, all *Caudoviricetes* vOTUs fell within clades also containing viralRefSeq representatives (**Figure 1c**). Thus, results indicate that *Caudoviricetes* vOTUs identified across the Waiwera estuary likely fall within already known *Caudoviricetes* viral orders or families. Metagenomics studies over the last decade have rapidly expanded our understanding of the diversity of DNA viruses in diverse habitats. While the findings here suggest that considerable aquatic DNA viral diversity remains to be discovered towards the tips of lineages (i.e., 47% of vOTUs in this dataset may represent novel genera), novel discovery at higher taxonomic ranks may be approaching an asymptote (i.e., most are *Caudoviricetes*, and likely fall within families or orders already characterised). The same saturation discovered here in aquatic systems may not be true of other habitats, however, which remain less studied and underrepresented in databases.

### Viral spatial distributions are constrained by the same habitat boundaries as putative hosts

Salinity and planktonic/benthic divides are major environmental barriers for the distribution of estuarine microbial communities^15^. There are also substantial differences in resource availability between water and sediment, and across non-saline, brackish, and marine zones that likely contribute to differences in community composition^15^. We previously showed that water, sediment, nitrite, and salinity are the major drivers of microbial (prokaryotic and eukaryotic) community composition differences within the Waiwera estuary^15^. Results here further suggest that microbial and viral community distributions are both partitioned according to the same major environmental parameters (**Figure 2; Figure S2**). Both prokaryotic and viral diversity was higher in saline than non-saline water and sediment samples (**Figure S1d**). In weighted beta diversity analyses for MAGs and vOTUs, salinity and sample type (water or sediment) each significantly explained ∼30% and ∼25% of the variability observed, respectively (salinity and planktonic versus benthic interaction *R*^2^=0.156 for MAGs and 0.138 for vOTUs; *p*-values=0.001) (**Figure 2a,b**). Viral communities in sediment and overlying water samples from the same location were more similar to each other when considering presence/absence alone (unweighted Bray Curtis dissimilarity), suggesting some connectivity between these two environments (**Figure S2a**). Despite often observing the same vOTUs in both sediment and overlying water samples, transcription was more clearly delineated by the environmental boundary (**Figure 2c; Figure S2b**). A similar pattern was observed across the non-saline/saline boundary in water. Together, these patterns highlight stark boundaries of viral activity between habitat zones (non-saline/saline and water/sediment). Less difference in virus distributions and activity was evident between brackish and marine zones in water or sediment. This likely reflects the greater overlap in the prokaryote composition of brackish and marine sites compared with freshwater sites^15^ (**Figure 2b**), possibly due to a greater similarity in osmoadaptive traits^42^.

**Figure 2.**
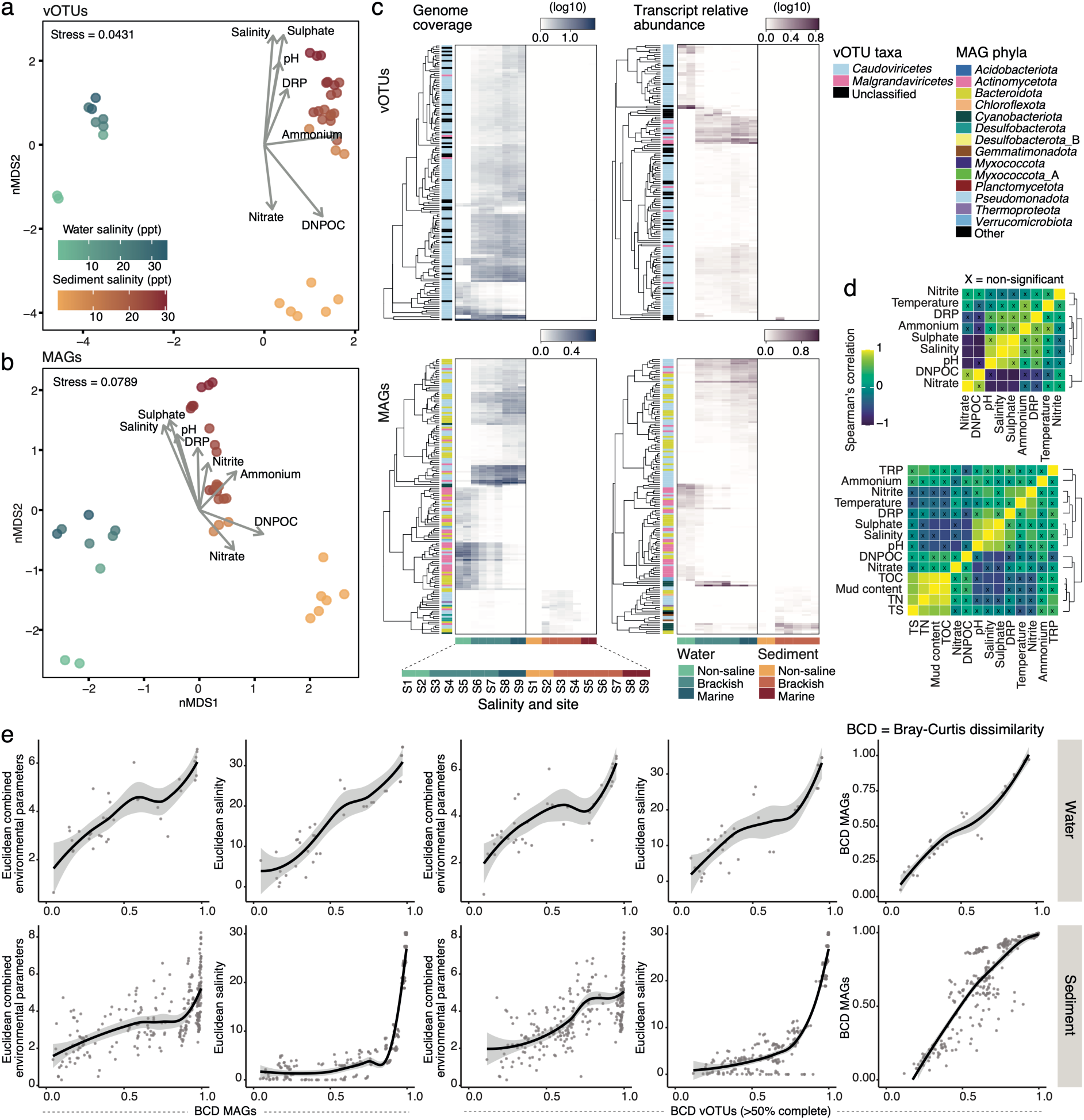
vOTU and prokaryote MAG distributions throughout the Waiwera estuary. nMDS ordinations of inter-sample comparisons based on weighted Bray-Curtis dissimilarities with fitting of environmental parameters for (a) vOTUs and (b) MAGs. (c) Heatmaps presenting the relative abundance (based on genome coverage) and transcriptional activity of the 150 most abundant and transcriptionally active vOTUs and MAGs (rows) by sample (columns). Numbers of vOTUs per estuary zone (non-saline, brackish, and marine) were 184, 109, and 494 for water, and 10, 24, and 36 for sediment, respectively. Numbers of MAGs per estuary zone were 84, 14, and 86 for water, and 55, 252, and 155 for sediment. (d) Correlation analyses of environmental variables measured in water (top panel) and sediment (bottom panel) samples. ‘X’ = non-significant (false-discovery rate-adjusted *p*-values >0.05). (e) Correlations between changes in vOTU and MAG community compositions (based on Bray-Curtis dissimilarities [BCD]) and environmental variables (based on Euclidean distance) in water (top row) and sediment (bottom row). Fitted polynomial regression lines are based on loess smoothing with shaded areas representing 95% confidence intervals.

We assessed the interacting effects of prokaryote and viral community compositions and environmental parameters (pH, salinity, ammonium, nitrate, nitrite, sulfate, dissolved reactive phosphorus [DRP], and DNPOC; for values see reference^15^). Among the measured environmental parameters, Spearman’s correlations showed that salinity co-varied strongly with sulfate and pH, which were higher in seawater, and with nitrate and dissolved non-purgeable organic carbon (DNPOC), which were higher in freshwater (**Figure 2d**). There was a strong correlation between prokaryotic (MAG) and viral (vOTU) Bray-Curtis dissimilarities across samples (Mantel test *r ≥*0.95, *p*-values=0.001 for water or sediment) (**Figure 2e**, **Table S5**). There was also a strong correlation between vOTU community dissimilarities and change in combined environmental parameters (Euclidean distance matrix; Mantel test *r* ≥0.74, *p*-values ≤0.002), as well as when testing salinity concentrations individually against viruses (*r* ≥0.84, *p*-values=0.001) or prokaryotes (*r* ≥0.89, *p*-values=0.001). Correlations between combined environmental variables or salinity and vOTU composition were no longer apparent in water samples after taking the effect of MAG composition into account (partial 3-way Mantel test, *p*-values ≥0.65), reinforcing the importance of host community composition on viral distributions. Although, the effect remained for sediment samples (*r*=0.30 for combined and 0.65 for salinity, *p*-values <0.01).

The strong coupling between microbial and DNA viral community composition patterns observed here has been seen, for example, in studies of agricultural soil^40^, the ocean over time and depth layers^29^, oxygen-deficient ocean water^43^, the global oceans^27^, and groundwater^37^. For open ocean and groundwater viruses, it has been suggested that viral community composition and distribution patterns are likely constrained by the distributions of hosts, which are in turn strongly influenced by environmental parameters that partition them into distinct ecological zones^27,37^. Phage host range can be highly specific and may be restricted to individual species or strains^44^. While phage adaptation can modify preferred host ranges^45^, this is likely limited to closely related hosts with similar routes to infection and defence repertoires. Alternatively, environmental parameters may directly limit the viability or infection success of DNA viruses. Phage infectivity and/or release of progeny have been shown to be directly affected by environmental conditions such as temperature^46^, pH^47^, and salinity^48–51^. For example, changes in salinity can result in the inactivation of viruses infecting extreme halophiles^48^ and marine prokaryotes over salt concentration gradients typical of estuaries^50^. Viral endolysins, important for lytic activity and the release of phage progeny, can also be inhibited by salt concentrations^51^. Constraints on estuarine viral distributions across habitat boundaries could therefore be influenced by the effects of physicochemistry, as explored further below, in addition to host range.

### Diversification within viral genera is constrained by ecological niche partitioning

Recent evidence suggests that viral phylogeny is delineated by environment type at low taxonomic ranks (e.g. species)^30^, implying viral adaptation to distinct ecosystems. In particular, viral phylogeny was shown to differ between terrestrial versus aquatic environments, with few viral clusters sharing sequences from both sources^30^. In light of this, and given habitat-associated differences in the distributions of estuarine viruses (**Figure 2a,b; Figure S2**), we examined the potential influence of ecological boundaries on viral diversification using clustering based on a protein-sharing network with high-quality environmental IMG/VR sequences, and inferred phylogeny via alignment of putative core genes. Of the 280 protein-sharing clusters that contained at least one vOTU in this study, around half contained sequences from exclusively non-saline (8%) or saline (50%) samples (**Figure 3a-c**). Within the remaining mixed-ecosystem clusters, water-column viruses were still predominantly skewed towards either non-saline or saline locations (**Figure S3a**). Some clusters with both non-saline and saline viruses may be due to misclassification of allochthonous viruses, such as may occur with viruses sampled from near-shore environments.

**Figure 3.**
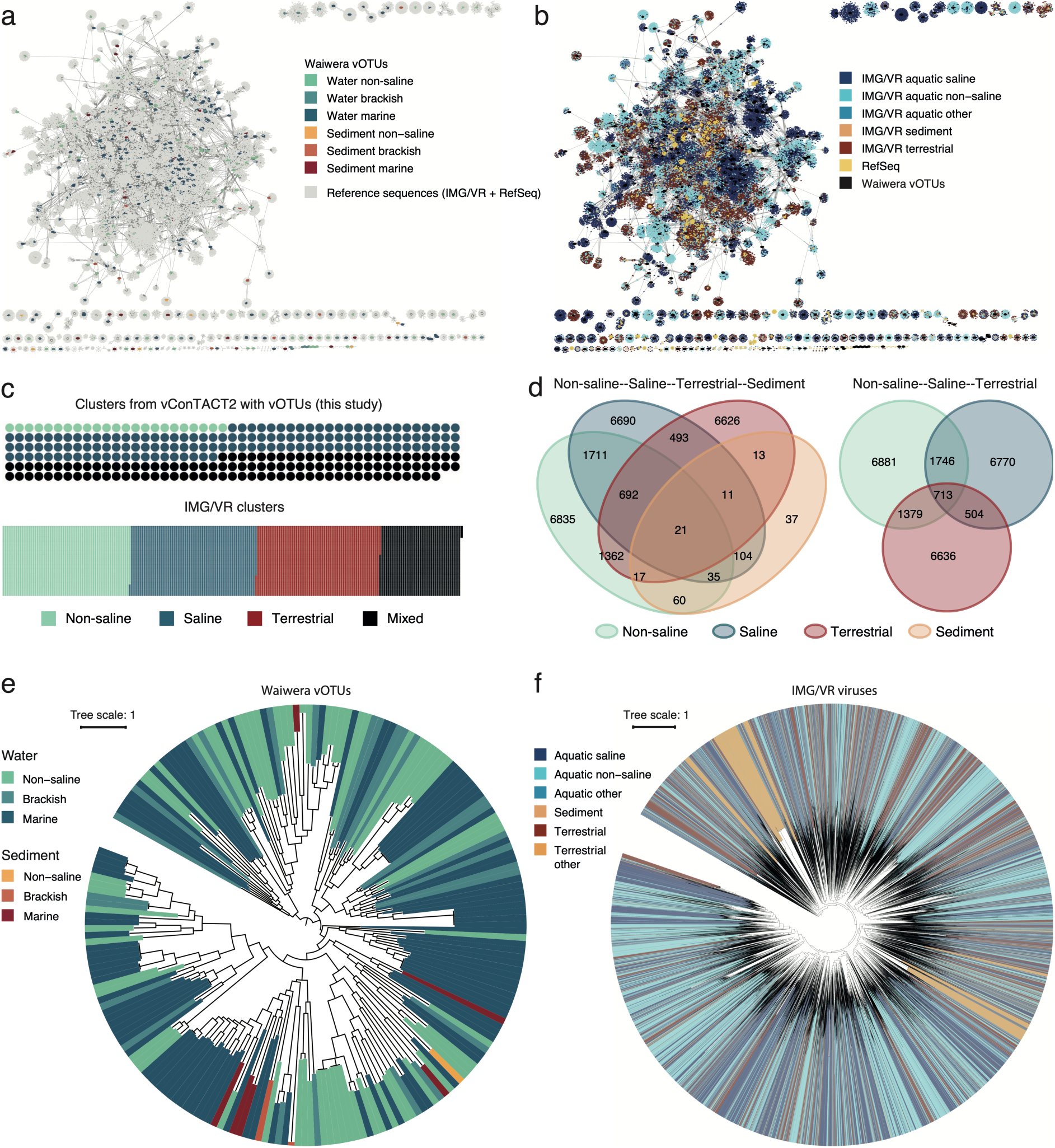
Viral diversification and ecological niche partitioning. (a-b) vConTACT2 protein-sharing networks incorporating Waiwera vOTUs (858 vOTUs >50% complete), viralRefSeq (v211), and all IMG/VR high-quality viruses that shared an interaction with a vOTU, coloured by (a) estuary ecosystem type (this study) and (b) IMG/VR reference ecosystem type. (c) Spot plot representation of individual clusters within the vConTACT2 network that included viruses from unique ecosystem types (non-saline, saline, terrestrial) or those with mixed ecosystem types for all clusters containing a Waiwera vOTU (858 vOTUs >50% complete) (top figure) and all clusters based on all high-quality IMG/VR viruses (bottom figure). (d) Venn diagrams depicting IMG/VR ecosystem types present in each vConTACT2 cluster (counts = number of clusters that fall into each region of the Venn diagram). (e) Inferred phylogeny for *Caudoviricetes* Waiwera vOTUs (205 vOTUs >85% complete) (lines extending from branch tips are coloured by estuary zone), and (f) IMG/VR high-quality viruses based on a concatenated protein alignment of predicted *Caudoviricetes* core genes (*n*=44) (lines extending from branch tips for are coloured by habitat). Trees were generated via IQ-TREE using ModelFinder (with VT+F+I+G4 selected). Scale bars represent amino acid substitutions per site. Numbers of Waiwera vOTUs per ecosystem type (a, b, c) are: 184, 109, and 494 for non-saline to marine water, 10, 24, and 36 for non-saline to marine sediment. Numbers of vOTUs per ecosystem type (e) are: 68, 26, and 102 for non-saline to marine water, 1, 2, and 6 for non-saline to marine sediment.

Extending this analysis to all clusters containing high-quality IMG/VR sequences from aquatic non-saline, aquatic saline, sediment, and terrestrial sources (including 288,511 IMG/VR sequences across 24,629 clusters) showed that most sediment-associated viruses clustered together with aquatic non-saline, aquatic saline, and/or terrestrial viruses (**Figure 3d**). Aqueous sediment environments likely consist of a complex mix of microenvironments, and may receive deposits of viruses from both water and terrestrial sources. When only considering aquatic non-saline, aquatic saline, and terrestrial viruses, 82.4% of clusters contained viral sequences from only one ecosystem type, and these were evenly distributed across the three habitats (aquatic non-saline=27.9% (6,881 clusters); aquatic saline=27.5% (6,770 clusters); terrestrial=26.9% (6,636 clusters)), with only 17.6% (4,342 clusters) containing viruses from mixed environment types (**Figure 3c-d**). This is in line with the strong effects that salinity and substrate/environment type exert on prokaryotic community composition. Clusters with mixed environment types were once again often dominated by viruses from either aquatic non-saline or aquatic saline sources (**Figure S3a**). To test the influence of potentially allochthonous viruses, viruses from “edge” environments (for example, intertidal zone, coastal, harbour, etc.) were excluded, resulting in only 530 clusters (2.2%; including 5346 viruses (1.9%)) containing viruses from both non-saline and saline environments. If clustering is taken to approximate distinct viral genera^41^, these data indicate that many viral lineages at approximately the rank of genus are adapted to and likely constrained by ecosystem zones based on salinity (i.e. aquatic non-saline vs. aquatic saline), while additional environmental factors likely drive viral niche partitioning between aquatic and terrestrial environments.

To further examine niche partitioning, phylogenetic relationships within the *Caudoviricetes* (representing 73% of estuarine vOTUs) were inferred via alignment of putative core genes. Several studies have attempted to infer phylogenetic relationships within the *Caudoviricetes* class based on trees of one or more conserved genes. For example, a terminase large subunit (TerL) protein tree suggested poor representation of soil DNA-virus diversity in viralRefSeq^30^, and demonstrated the potential utility of TerL for discriminating major differences among *Caudoviricetes*. However, there and in this study, TerL trees were inconsistent with RefSeq viral taxonomy, with viral families separated over distinct clades (**Figure S3b**). Concatenated protein alignments based on multiple single-copy core genes represents an alternative. Work highlighting the polyphyletic nature of DNA virus taxonomy (prior to recent taxonomy revisions^52^) inferred *Caudoviricetes* phylogeny via trees generated from concatenated protein alignments (based on hits with the VOG database)^53^. For viralRefSeq *Caudoviricetes* genomes, there was overall agreement between the tree generated using this method and the current viralRefSeq taxonomic affiliations (**Figure 1c**). The environment types associated with vOTUs from this study (based on where they were most abundant) were interspersed throughout the trees, indicating that at high taxonomic ranks within the *Caudoviricetes*, members are present across environmental divides (**Figure 3e**). However, most sub-clades of closely related viruses were again only associated with one ecological niche type, supporting protein-sharing network analyses here and previously^30^. Similar results were obtained when using all high-quality aquatic and terrestrial *Caudoviricetes* viruses from the IMG/VR database (*n*=128,266) (**Figure 3f**).

In the IMG/VR analyses, only 20% (23,754) of included genomes met the filtering criteria when using the core genes identified based on *Caudoviricetes* viruses in viralRefSeq. As highlighted by work on soil viruses^30^, viralRefSeq does not yet include representatives covering the full genomic diversity of *Caudoviricetes* viruses globally, and core genes selected from viralRefSeq alone may not be appropriate for all *Caudoviricetes* virus lineages. When putative core genes were re-identified based on all IMG/VR *Caudoviricetes*, a smaller set were identified (13 versus 44) (**Table S6**), but the resulting concatenated protein alignment retained a much greater number of viruses (85,588) (**Figure S3c**). In either case, the overarching patterns for ecosystem associations remained the same. The finding that fewer “core genes” met the filtering criteria when core genes were identified from all IMG/VR viruses identified as *Caudoviricetes*, indicates a complex and diverse phylogeny within IMG/VR *Caudoviricetes* viruses. Due to their genomic diversity, the true phylogeny of global *Caudoviricetes* viruses (as well as DNA viruses from other classes) may not be adequately captured by a single set of core genes. In future, a decision tree approach using a nested hierarchy of sets of core genes that delineate key branching events and relationships within subclades may better describe the complex phylogeny of DNA viruses globally.

In summary, multiple lines of evidence support the finding that ecological niche partitioning is evident for DNA viruses. Representatives within high taxonomic ranks (approximately family level and above) are found across ecological zones, suggesting that DNA viruses often evolved agnostic of ecological context at least as far as family level diversification. However, at lower ranks (approximating genus level), virus diversification is largely partitioned by ecological niche based, at least indirectly, on factors such as salinity and the aquatic-terrestrial divide. Similar observations have been made of prokaryotes, where, for example, members of individual genera or species are rarely found across freshwater-saline boundaries, although taxa on either side of this boundary are often related at approximately family level^22^. As viruses tend to be host range limited^27,44^, it is not unexpected that they show similar partitioning. Although the data we analysed represents a considerable resource of high-quality viral genomes (IMG/VR database *n* = 288,511; estuarine database *n* = 858), the global virome remains under-sampled. As reference databases continue to expand, additional studies are warranted to further examine the extent to which closely related viral taxa are partitioned into ecological niches.

### Molecular signatures of environmental adaptation in DNA viruses

#### Changes in viral gene amino acid encoding associated with environmental salinity

Osmoadaptations can confine prokaryotes to a set niche, resulting in transitions between freshwater and saline environments that are rare and evolutionarily ancient^22,23^. Genomic signatures of adaptation to saline environments are well described for prokaryotes and include adaptations necessary to maintain protein stability^19,20^. In halophiles, acidic and basic amino acids are disproportionately found at the exposed surfaces of proteins^54^. Excess acidic amino acid residues on protein surfaces enhance protein solubility and structural rigidity, and promote the establishment of a hydration shell, all of which are thought to protect against the denaturing effects of high salinity^42,54–56^. Alterations in amino acid encoding to increase protein stability are a common feature of salt-adapted prokaryotes, and occur irrespective of phylogeny, indicating convergent adaptation to high salt environments^20,54,56,57^. More recently, this trait was observed at estuarine salinities^58^ and salinities comparable to those of estuaries^22,42^. At these salinities, protein acidity is most pronounced in the secreted proteins of prokaryotes^42^, consistent with use of the salt-out approach^15,59^, and higher extracellular versus intracellular salinities.

We sought to examine whether similar genomic signatures of environmental adaptation were also present in viral genomes, reflecting adoption of the same approach to stabilise proteins exposed to saline conditions. Based on findings for prokaryotes^42^, we further hypothesised that the selective pressures of environmental adaptation may be stronger on proteins that must be functional in the conditions of the surrounding environment (e.g. tail, capsid, and spike proteins) or in intracellular zones that may be influenced by external conditions (e.g. endolysin proteins), compared with those that function within the host cell without exposure to external conditions (e.g. large terminase and DNA polymerase proteins). First, we examined patterns of encoded amino acid proportions in predicted viral genes across the Waiwera estuary to directly analyse changes in genomic signatures against gradual increases in salinity. We then analysed all high-quality IMG/VR viral genomes (*n*=492,846) to investigate whether the same patterns were reflected in viruses from diverse ecosystems.

In the estuarine dataset the overall ratio of acidic (glutamic and aspartic acid) to basic amino acids significantly increased with increasing salinity in all predicted protein groups analysed (*p*-values ≤0.001) except DNA polymerase (*p*=0.213). The trend was strongest for major tail, major capsid, and endolysin proteins (*R*^2^ values = 0.21, 0.03, and 0.05, respectively) compared to minimal change in tail tape measure, large terminase (TerL), or DNA polymerase proteins, or the predicted proteome as a whole (*R*^2^ values ≤0.01) (**Figure 4a; Figure S4a**). In linear models comparing only viral proteins from saline samples we found that the effect sizes were reduced for major capsid and endolysin proteins (albeit significant, *R*^2^ <0.01, *p*-values <0.02), and the trend was lost for major tail proteins (*R*^2^ <0.01, *p*-values *=* 0.372). This suggests there may be a threshold effect at the non-saline-to-saline boundary. The observed increase in ratios of acidic-to-basic amino acids with increased salinity, across this threshold, is small. However, charged amino acids are predominantly found at the exposed surfaces of proteins rather than on unexposed surfaces or buried within protein interiors^54^, therefore it is not unexpected that effect sizes are small when considering the protein as a whole. Assuming the same trends in viruses as in prokaryotes, the variability observed within samples is also likely to be due, in part, to the influence of lineage-specific biases on osmoadaptive trends. Differences exist in the amino acid osmoadaptations among prokaryotic taxa^60–62^.

**Figure 4.**
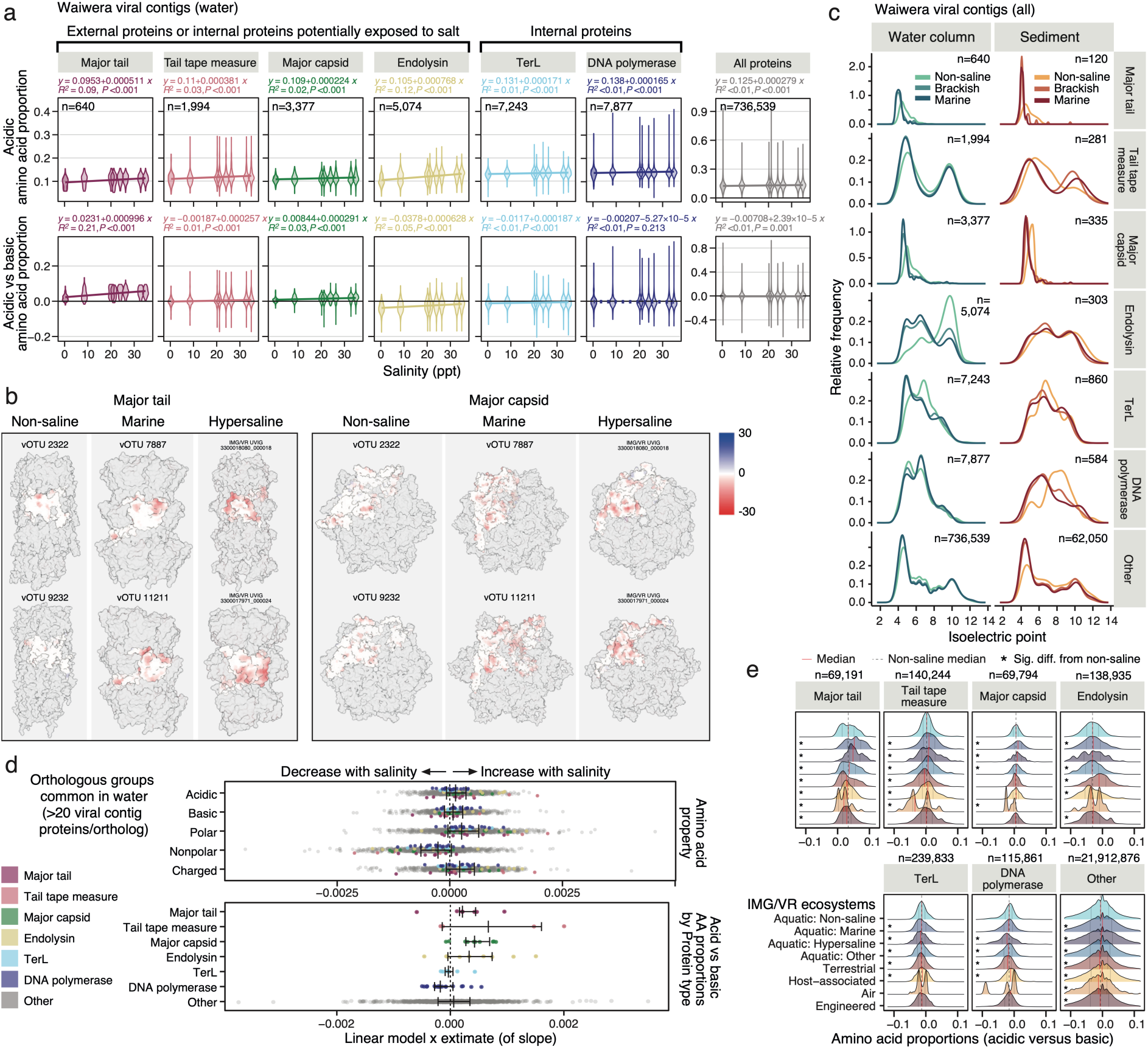
Molecular signatures of environmental adaptation in DNA viruses. (a) Proportions of acidic amino acids (top row), and the difference between the proportion of acidic-basic amino acids (bottom row) in predicted viral contig (*n*=70,326) protein subsets and all combined proteins in the estuary water column. Linear modelling equation and statistics were calculated via the *stat_poly_line()* and *stat_poly_eq()* functions from the *ggpmisc* package in R, with line shading representing 95% confidence interval. Line is solid (versus dashed) for significant correlations. (b) Representative protein complex structural predictions and surface electrostatic potentials (ESP; blue to red surface coloring) for major tail protein (left) and major capsid protein (right) for two representatives each from non-saline, marine, and hypersaline environments. Protein structures were predicted from protein sequences via Alphafold 3 server, with visualisation and ESP calculation in ChimeraX. (c) Predicted protein isoelectric points by estuary salinity zone. (d) x-estimates from linear models showing the trend of changes in amino acid proportions (categorised as: acidic, basic, polar, non-polar, and charged) with respect to increases in salinity across the estuary in water, with bars representing median and interquartile range. Plots represent orthologous groups per protein type. Numbers of orthologous groups (and sequences) per protein type: major tail = 7 (364); tail tape measure = 4 (201); major capsid =10 (3,272); endolysin = 9 (2,131); TerL = 7 (2,874); DNA polymerase = 17 (4,958); and total = 1,230 (184,695). (e) The difference between acidic and basic amino acid usage in predicted proteins subsets from all high-quality viruses in the IMG/VR database, categorised by environment type, with vertical lines representing median (red) and interquartile range (grey). Dotted vertical lines show the median for the Aquatic: Non-saline group. * = significant difference in pairwise comparisons (Dunn’s test of multiple comparisons with bonferroni correction) against the Aquatic: Non-saline group.

The stronger effect observed for major tail, major capsid, and endolysin proteins between non-saline and saline samples suggests an increase in acidity of the proteins that is most likely to be influenced by the external environment, and which does not solely reflect biases inherited due to reliance on host machinery (see discussion below). Spike proteins represented an exception with no association between salinity and the ratio of acidic to basic amino acids detected (*p*=0.133). Spike proteins are under strong selective pressure based on their interaction with host receptors^34^, which may override other environmental selective pressures. Viral adaptations are likely constrained by a variety of factors, including host machinery^35^, host defense mechanisms^32,33^, protein-protein interactions (e.g. due to the effects of acidity and surface electrostatic potential)^63^, and structural limitations (e.g. physical constraints on capsid size, shape, stability, and protein complex interactions)^64,65^.

Amino acid-based observations were reflected by protein surface electrostatic potentials (representatives, **Figure 4b**; examples of 70 proteins, **Figure S5**) and protein isoelectric points (pI). Isoelectric points describe the pH at which surface charge changes between positive and negative, and were lower at higher salinities, particularly in major tail, major capsid, and endolysin proteins (linear model *R*^2^ values = 0.18, 0.02, and 0.07, respectively) (**Figure 4c; Figure S4b**). In addition to the role of protein acidity in maintaining protein stability^56^ and impacting protein-protein interactions^63^, the pI of viral particles also influences their colloidal behaviour and sorption, and this is largely determined by the pI of outer proteins such as coat or envelope proteins^66–68^. In our study, predicted major tail and major capsid proteins from all estuary zones had a narrow range of low pI relative to the rest of the viral proteome (**Figure 4c**). Despite this skewed baseline, statistically significant reductions in the pI of these proteins were observed with increasing salinity across the estuary (*p*-values <0.001) (**Figure S4b**). The influence of pI on the colloidal behaviour and sorption of viral particles, protein-protein interactions, and the need to maintain adequate protein function in the conditions of the external environment likely represent interacting selective pressures driving viral adaptation based on environmental salinity.

To identify whether the trend above is influenced by distinct protein compositions across the salinity gradient, individual viral orthologous groups (VOGs) that spanned the estuary were inspected. As salinity increased, the proteins from each of these VOGs tended to have increased proportions of acidic, basic, and polar amino acids, and reduced nonpolar amino acid proportions, indicating that there is selective pressure to adapt amino acids within protein orthologs (**Figure 4d; Figure S4c**). There was also a consistent trend for reductions in amino acids with codons more often containing G and C nucleic acids (e.g. alanine, proline, glycine, arginine, tryptophan; **Figure S4d**). This is consistent with our observations of reduced genomic GC content in aquatic viruses from saline environments (see below). An exception was for the two acidic amino acids (glutamic and aspartic acid), which also have a moderate rate of G and C nucleic acids in their codons. This suggests that the selective pressures driving acidic amino acid enrichment and genomic GC content operate independently.

Using IMG/VR high-quality genomes, we likewise observed that major tail, tail tape measure, and major capsid – but not endolysin, large terminase, or DNA polymerase – proteins from aquatic saline and hypersaline viruses had significantly more acidic proteins (acidic vs basic amino acid proportions) than those from aquatic non-saline and terrestrial sources (Dunn’s test adjusted *p*-values <0.001), and that differences were more pronounced for these proteins than for the proteome as a whole (**Figure 4e; Table S7b**). This trend is most obvious among the *Duplodnaviria* (which contains the class *Caudoviricetes*) (**Figure S6a**). This realm dominates the IMG/VR dataset (72% of high-quality viral sequences), and most of the tested protein categories (e.g., major tail, tail tape measure, large terminase) were rarely identified in the other realms. As per the estuarine dataset, these patterns were evident in the relative shifts of protein pI when comparing between aquatic non-saline, aquatic saline, aquatic hypersaline, and terrestrial viruses (**Figure S6b**). Finally, for both the estuarine and IMG/VR datasets, the proteome of viruses from saline sources also had greater proportions of polar versus non-polar amino acids. The latter was primarily driven by a marked reduction in proportions of alanine (**Figure 4d; Figure S4a,c**), as is also found in prokaryotes^22^. A single base substitution in the middle position modifies codons for alanine to instead code for the acidic amino acids glutamic or aspartic acid. Alanine is thought to have a relatively neutral effect on protein stability^22,69^, which may explain the conserved adaptation of substituting alanine for acid amino acids in saline environments. Overall, our observations indicate that adaptations to salinity, common to prokaryotes, are also present in viruses.

#### Proportions of viral gene GC content and environmental niche

Based on Waiwera estuary data, viral gene GC proportions decreased linearly and significantly with increasing salinity in water and sediment. In water samples, this pattern was observed consistently for all genes combined and each gene category (major tail, tail tape measure, major capsid, endolysin, large terminase, and DNA polymerase) (fitted linear model *R*^2^ values = 0.06-0.20; *p*-values <0.001) (**Figure 5a**), as well as in linear modelling of individual VOGs present across the estuary water column (**Figure S4e**). The factor driving differences in viral gene GC content across the estuary appears to act on the whole genome, independent of the cellular location of encoded proteins and potential salt exposure, unlike in trends observed for amino acids. This again suggests that salinity was not a driver of viral GC content trends in the estuary. Moreover, elevated, rather than reduced, proportions of genomic GC content are often associated with prokaryotes from hypersaline environments^20,70^. This is not ubiquitous, and key traits of salinity adaptation, including amino acid biases, are present irrespective of overall GC content, which may co-vary due to other associated environmental factors such as nitrogen limitation^71^ or UV-induced damage^20,70^. As such, increased overall GC content previously observed in halophiles may be a factor of the environments that many halophiles commonly inhabit, independent of salinity.

**Figure 5.**
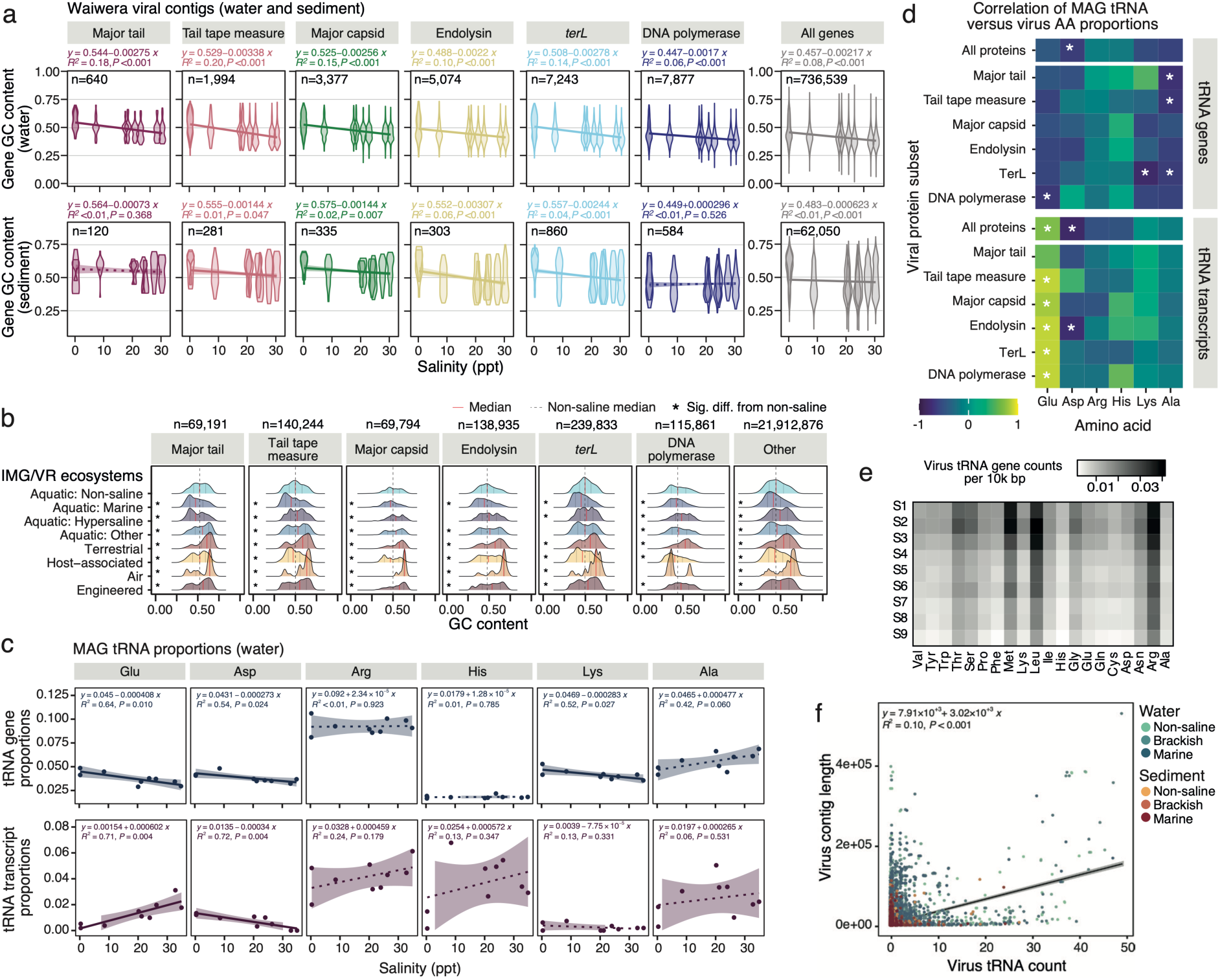
(a) GC content of genes per functional category in water (top row) and sediment (bottom row). (b) Viral gene GC content by environment type for all high-quality IMG/VR viruses, split by gene functional category, with vertical lines representing median (red) and interquartile range (grey). Dotted vertical lines show the median for the Aquatic: Non-saline group. * = significant difference in pairwise comparisons (Dunn’s test of multiple comparisons with bonferroni correction) against the Aquatic: Non-saline group. (c) Total proportions of prokaryote tRNA gene (top row) and transcript (bottom row) abundances against salinity in the Waiwera estuary (water) for acidic (glutamic and aspartic acids) and basic (arginine, histidine, lysine) amino acids and alanine. (d) Spearman’s correlation analyses comparing predicted virus protein amino acid and cognate prokaryote tRNA gene (top panels) and transcript (bottom panels) relative proportions across the Waiwera estuary water samples for virus protein subsets and all proteins combined. * = Spearman’s *p*-value ≤0.05. (e) Heatmap of virus tRNA gene counts per 10k bp split by amino acid. (f) Correlation of virus contig lengths against virus tRNA gene count per contig. Linear modelling equations and statistics (a, c, f) were calculated via the *stat_poly_line()* and *stat_poly_eq()* functions from the *ggpmisc* package in R, with line shading representing 95% confidence interval. Line is solid (versus dashed) for significant correlations.

Analysis of IMG/VR viruses further showed proportions of viral gene GC content differed strongly based on the virus source environment, indicating that factors influencing viral genomic GC content are broad (**Figure 5b; Figure S7a**). Viruses from aquatic marine sources had significantly reduced proportions of GC content overall compared to aquatic non-saline viruses (respective medians and interquartile range 0.416 and 0.346-0.502 versus 0.448 and 0.367-0.536) (Dunn’s test of multiple comparisons *p*-values <0.00001; **Table S7c**), reflecting findings from the Waiwera estuary. Viral genomes from aquatic hypersaline environments and terrestrial sources (irrespective of salinity) were more GC-enriched (respective medians and IQR: 0.495 and 0.406-0.582; 0.538 and 0.438-0.611.

Dunn’s test of multiple comparisons *p*-values <0.00001). Elevated GC content is also seen in prokaryotes from terrestrial environments compared to ocean surface water environments^72^. GC content distributions for viruses from wetlands, groundwater, sediment, deep subsurface, plant litter, and the air were multimodal and tended to span the ranges of GC proportions characteristic of aquatic and terrestrial viruses (**Figure S7a**), and may reflect distinct microenvironments within these ecosystems. When subsetting by viral realm, these differences when comparing between viruses from aquatic non-saline, aquatic marine, hypersaline, and terrestrial environments were more strongly associated with *Duplodnaviria, Monodnaviria*, and *Varidnaviria* viruses than with *Riboviria* (**Figure S7b**), suggesting that underlying differences in viral biology or host domain between the realms influence the association between virus GC content and environmental source. For example, the link between virus and host GC content and codon usage bias is stronger for bacteriophage than for eukaryotic viruses^35^.

It remains unclear whether the observed associations between viral GC content and environment types are a result of direct environmental adaptation of viruses themselves. A recent study examining viral genomic content in relation to broad host categories for ∼10,000 viral genomes also found multi-modal viral GC contents, largely driven by a trimodal pattern for dsDNA viruses dominating the dataset^31^. Viral host characteristics may underlie this multi-modal distribution due to host-related selective pressures on viral genome content^31^, which can include host translational biases^73^ and viral-host gene exchange^74^. Our data show that virus GC content can be further differentiated based on environment type (such as aquatic non-saline, aquatic saline, aquatic hypersaline, and terrestrial).

However, we cannot discount that this may be an indirect effect driven by environmental adaptation of the host. Selective pressures acting directly on host GC content may be acquired by viral genomes via host-specific adaptation. Alternatively, direct selective pressures related to both the host and environment type could work in parallel to determine the GC content signatures of phage.

### Virus amino acid proportions and changes in prokaryote translational machinery

Virus protein translation largely relies on the pool of host tRNAs^35^, and the relationship with tRNA pools and codon usage bias (CUB) in genes is strongest for highly expressed genes^75^. To assess whether the observed changes in amino acid biases in virus proteins are driven by underlying changes in prokaryote (i.e., putative host) machinery, rather that direct environmental adaptative pressure, we investigated (a) changes in MAG tRNA repertoires across the estuary (and influence of phylogeny), (b) associations between MAG tRNA gene or transcript proportions and amino acid proportions in predicted viral proteins, and (c) the presence of additional endogenous tRNA genes in virus genomes. UniFrac analysis of concatenated alignments of prokaryote core genes (derived from GTDB-Tk) revealed graduated shifts in phylogeny across water (non-saline to marine) and sediment (brackish to marine) samples (**Figure S8a**). However, prokaryote tRNA repertoires (based on Bray-Curtis dissimilarities of tRNA counts per amino acid) did not cluster according to phyla or estuary habitat type (**Figure S8b**). Likewise, when considering actinobacterial Luna cluster genera, which were relatively abundant in either non-saline (*Rhodoluna*) or saline (*Aquiluna*) water samples, tRNA counts for each amino acid did not cluster by lineage or estuary habitat zone (non-saline, brackish, or marine) (**Figure S8c**).

When comparing prokaryote tRNA repertoires and virus amino acid encoding, we found that virus amino acid proportions for glutamic acid, which increased significantly with increasing salinity (linear model *p*-values <0.005) (**Figure S10**), also significantly positively correlated with cognate prokaryote tRNA gene transcript frequencies (significant correlations for all viral proteins combined, as well as all protein subsets except major tail protein; *p*-values <0.05) (**Figure 5c-d**). This indicates that host-adaptation could be a primary driver of the predicted changes in glutamic acid proportions in virus proteins in this study, and may have contributed to the small shift in overall acidity observed with salinity in the viral proteins as a whole (**Figure 4a**). The reduced association observed for major tail proteins (**Figure 5d**) suggests that additional factors beyond host-adaptation also influence the acidity of these proteins in particular.

With the exception of glutamic acid, however, no significant positive correlations were observed between acidic or basic amino acid proportions in predicted virus proteins and prokaryote tRNA genes or transcripts for these amino acids when analysing all virus proteins together or for protein subsets (**Figure 5d; Figure S9c**). For example, frequencies of tRNA genes for both acidic amino acids and the basic amino acid lysine significantly reduced with increasing salinity across the Waiwera estuary (*p*-values <0.05) (**Figure 5c**), contrasting with the changes in amino acid usage in virus predicted proteins (**Figure S9a**). While virus protein proportions of aspartic acid increased with increasing salinity (*p*-value <0.001), these increases were significantly negatively correlated with proportions of cognate tRNA genes and transcripts, which significantly reduced with increasing salinity (*p*-values <0.05) (correlation for all proteins, Spearman’s Rho values <-0.65; *p*-values <0.05) (**Figure 5d; Figure S9c**). Therefore, the relative encoding of aspartic acid appears to be decoupled from adaptations based on putative host translational machinery, and may represent a flexible, independent mechanism for osmoadaptation.

We assessed whether differences in the carriage of additional tRNA genes in virus genomes may be one mechanism enabling viruses to modulate amino acid proportions independent of host adaptation. tRNA genes in Waiwera viral contigs ranged from 0-49 (overall mean per contig = 0.17), with tRNAs for the amino acids arginine, leucine, and methionine being the most prevalent regardless of sample site (**Figure 5e**). A small but significant reduction in the total number of virus tRNA genes per genome was observed with increasing salinity across the estuary (mean of means for non-saline [sites 1 and 2] = 0.32/genome, brackish [sites 3-7] = 0.20/genome, and marine (sites 8 and 9) = 0.12/genome; linear model *R*^2^ <0.01, *p* <0.001), which may, at least in part, be related to differences in average genome size recovered. Virus contig sizes similarly reduced with increasing salinity (mean of means for non-saline = 9112 bp, brackish = 8987 bp, and marine = 8343 bp; linear model *R*^2^ <0.01, *p* <0.001), and virus tRNA gene counts had a moderate positive relationship with virus contig size overall (linear model *R*^2^ = 0.10, *p* <0.001) (**Figure 5f**). When assessing changes in the relative abundances of tRNA genes for each amino acid, proportions of tRNAs for the two acidic amino acids (aspartic and glutamic acid) and the basic amino acids histidine and lysine all significantly decreased with increasing salinity, while tRNAs for arginine significantly increased (linear model *p*-values <0.01). The same trend was seen with proportions of prokaryote tRNA genes for aspartic acid, glutamic acid, and lysine (**Figure 5c**), suggesting that relative proportions of tRNA types in virus genomes may be determined by that of their hosts, and indicating that differential virus carriage of tRNAs is unlikely to be the mechanism enabling protein adaptation in response to increased salinity.

The relationship between host tRNA pools and codon usage bias (CUB) for individual genes in both host and virus genomes is complex. Translational biases are not consistent across different virus functional gene types: structural proteins are more adapted to host CUB than non-structural virus proteins^73,76^, while the selective pressure to match CUB is minimal in the rest of the virus proteome^73^. Differences in relative codon usage biases have also been observed for virus genes involved in early vs late stage of infection^77,78^, across different host tissues within individual eukaryote hosts^79^, and in generalist vs specialist viruses (with respect to host range)^80^. CUB also differs within individual genes, where sub-optimal codon usage at the start of mRNA can increase overall efficiency of translation by reducing the effects of jams due to ribosome stacking^81^. Furthermore, while CUB similarity between virus and host can increase virus protein translation (due to availability of cognate tRNAs), this also has the trans-regulatory effect of differentially depleting the tRNA pool, impeding host translation^82^, and resulting in deleterious “symptomatic” infection states for the host cell. This trans-regulatory effect may drive a counterbalancing adaptive pressure against viruses matching host CUB^82^. Further to this, our results highlight a disconnect between prokaryote machinery (tRNA gene and transcript proportions) and virus amino acid encoding within sampled estuarine communities. However, our analyses were based on the tRNA repertoires and amino acid encoding of prokaryote and viral communities, which may obscure direct relationships between true virus and host pairs. Studies analysing patterns in clearly defined virus-host pairs are needed to further clarify the effects of the competing adaptive pressures of host machinery and environmental conditions.

### A rare case of closely related vOTUs spanning the salinity divide

Further analyses identified a rare example of closely related vOTUs in our estuary dataset that spanned the “salinity divide” between the non-saline and saline habitats. These vOTUs (vOTU_12982 and vOTU_19433), classified as *Caudoviricetes* (*Duplodnaviria* realm), were closely branching in the reconstructed core-gene phylogeny (**Figure 6a**) and clustered together in network analysis, suggesting a relatedness at approximately genus level or lower. However, the distributions of these two vOTUs were clearly differentiated into non-saline (vOTU_12982) and saline (vOTU_19433) zones based on genome and transcriptome abundances (**Figure 6b**). These vOTUs also clustered together with vOTU_5409 and vOTU_6591 (excluded from the “core genes” phylogeny due to low predicted completeness), which shared a similar “non-saline” distribution pattern as vOTU_12982.

**Figure 6.**
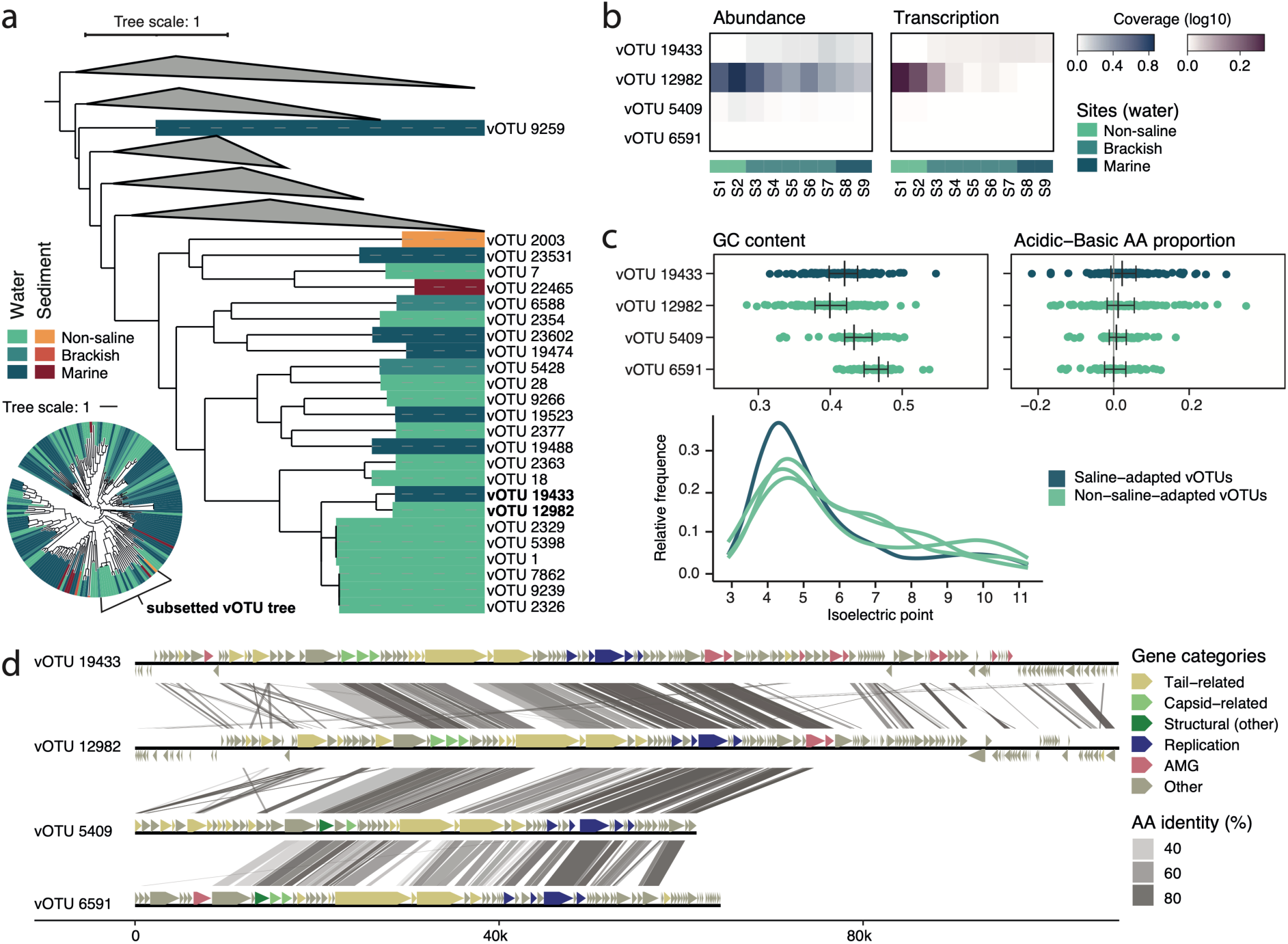
A comparison of vOTUs spanning the salinity divide. (a) Clade from the phylogeny for *Caudoviricetes* vOTUs (Figure 3e) showing closely related (short branching) vOTUs 19433 and 12982, which span the “salinity divide”. (b) Heatmaps of genome abundance and transcription across the estuary water column for these and two other closely related vOTUs. (c) Comparisons between gene GC content, the difference between encoding of acidic-basic amino acids, and protein isoelectric points for the four vOTUs, with bars in dot plots representing median and interquartile range. Each dot in the figure represents values for individual genes (left) or predicted proteins (right). (d) Gene synteny plot for the four vOTUs showing protein homology, including conserved and variable regions in the genomes.

Gene synteny analysis highlighted a suite of replication- and structure-related genes that are well conserved (∼80% AAI) across all four vOTUs, together with some highly variable “accessory” regions (**Figure 6d**). Reduced homology was observed among the tail-related genes of these vOTUs, which may reflect adaptation to distinct hosts. No reliable CRISPR spacer-based host matches were obtained for these vOTUs (**Supplementary Materials**). However, the likelihood of distinct hosts is supported by the variable genomic GC contents among the four related vOTUs (pairwise Dunn’s test with Bonferroni correction *p*-values <0.01) (**Figure 6c; Table S7d**), which was also evident when restricting analyses to genes in common for the two vOTUs with complete genome sequences (vOTUs 19433 and 12982 have circular genomes with identified direct terminal repeats) (**Figure S11a**). Finally, while the four vOTUs were predicted to be closely related, trends of amino acid encoding and pI of predicted proteins were consistent with our findings outlined above, with increased ratios of acidic to basic amino acid encoding and a shift to lower protein pI in the “saline” vOTU_19433 when compared to the three “non-saline” vOTUs (**Figure 6c; Table S7e**). Future studies could further explore the occurrence of closely related viruses across distinct habitats, and their underlying adaptive traits, using global datasets such as the IMG/VR database.

### Summary

We identified that environmental distributions and diversification within DNA viral genera are often constrained by ecological niche partitioning based on salinity and planktonic/benthic divides. This was observed both in our estuarine dataset, and based on an analysis of globally distributed viruses. Strong correlations were observed between estuarine viral and prokaryote community distributions suggestive of host range limitation. Additionally, our results indicate that genomic signatures of salinity adaptation are present in viruses independent of host-adaptation, highlighting the likely role of environmental salinity – together with other factors – in driving viral adaptation and niche differentiation. These include increased usage of acidic amino acids, elevated ratios of acidic to basic amino acids, reduced alanine contents, changes in protein surface electrostatic potentials, and decreased protein isoelectric point as salinity increases, particularly for those proteins that must maintain proper function in the external environment (for example, tail and capsid proteins). In some cases these changes were decoupled from adaptations based on host translational machinery (tRNA gene and transcript proportions), and may represent a flexible mechanism for osmoadaptation. While we cannot discount other factors influencing amino acid biases in the viral proteome, the modification of protein amino acid repertoire in response to environmental salinity appears to be a common adaptive strategy spanning both prokaryotes and viruses. These results have implications for our understanding of viral ecology and the limits of viral viable ranges. For example, recent work has highlighted the prevalence of cross-biome viral dispersal via the natural water cycle^83^. However, our findings indicate that successful proliferation of viruses transferred to distinct biomes (e.g. freshwater, saline, terrestrial) is limited, and is likely constrained by genomic adaptations tailored to specific environmental niches in addition to host range limitation.

## METHODS

### Data acquisition

#### Waiwera estuary metagenomics and metatranscriptomics

Metagenomic and metatranscriptomic Illumina sequencing was conducted on filtered water and triplicate sediment samples from nine sites in the Waiwera estuary (*n*=9 water samples; *n*=27 sediment samples with three spatial replicates per site) as described previously^15^ (see **Supplementary Materials** for further details; **Table S8**). Filtered water samples included viruses associated with cells or biofilms trapped on 0.22 micron filters. Samples spanned the salinity gradient (0 to 35 ppt) and large differences in nutrient availability^15^.

#### IMG/VR database

High-confidence genomes were downloaded from the IMG/VR database v7.1^84^, and filtered to retain sequences meeting MIUViG^85^ quality thresholds of ‘High-quality’ or ‘Reference’ (*n*=492,846 sequences). This dataset was used for vOTU taxonomy prediction, gene GC, amino acid proportion, isoelectric point analyses, and inference of *Caudoviricetes* phylogeny (see below). A subset of these data was generated by filtering for environmental viruses based on IMG/VR metadata “Ecosystem Classification” (*n*=288,511 sequences) for use in analyses of viral clustering and phylogeny based on environment types (see below).

### Data processing

#### RNA sequence processing

RNA sequencing reads for water samples (sites 1-9) and sediment samples (sites 1-7) were quality trimmed and filtered via Trimmomatic v0.39^86^ with the following parameters: ILLUMINACLIP:1:25:7 CROP:115 SLIDINGWINDOW:4:30 MINLEN:50. Residual rRNA sequences were removed using SortMeRNA (v4.3.6; database: smr_v4.3_fast_db)^87^ (**Table S8**). Sediment sites 8 and 9 were excluded from RNA sequencing due to low quality^15^.

#### Recovery of prokaryote metagenome-assembled genomes

Prokaryote metagenome-assembled genomes (MAGs) were recovered as previously described^58^. In brief, initial binning was conducted via CONCOCT (v0.4.1)^88^, MetaBAT (v2.12.1)^89^, and MaxBin (v2.2.4)^90^ using contig coverages. Coverages were determined based on reads mapped via BBMap v37.93^91^. MAGs were dereplicated across tools via DAS_Tool v1.1.1^92^. MAGs were then dereplicated across assemblies via dRep v1.4.3^93^ at a threshold of 98% average nucleotide identity (ANI), resulting in 646 unique MAGs (**Table S9**). MAG quality was further assessed with CheckM v1.2.1^94^, taxonomy predicted using GTDB-Tk (v2.4.0; database v214)^95^, and gene predictions and annotations generated via DRAM v1.3.5^96^.

#### Virus identification

Viral sequences were identified by VIBRANT v1.2.1^97^, VirSorter2 v2.2.3^98^, and deepVirFinder v1.0^99^, as previously described^37^, with the following modifications: For each assembly, putative viral contigs were dereplicated using a custom script (virome_per_sample_derep.py; https://github.com/GenomicsAotearoa/workflow.DNA_virus_niche_adaptations). Where prophages were identified by VIBRANT or VirSorter2, trimmed prophage sequences were retained. Clustered virus population units (henceforth, virus operational taxonomic units (vOTUs)^85^) were generated across assemblies based on ≥95% ANI over ≥85% coverage of the shorter sequence, via the Cluster_genomes_5.1.pl script (https://github.com/simroux/ClusterGenomes). vOTUs were assessed via CheckV v 0.7.0^100^, and filtered based on the following criteria: contig_length >3000 bp AND (viral_gene>0 OR (viral_gene==0 AND host_gene==0)). vOTU gene predictions and annotations were generated via DRAM-v v1.4.6^96^.

#### vOTU taxonomy

vOTU taxonomy was predicted via multiple approaches, including, in order of priority: 1. clustering based on a protein sharing network of vOTUs and viralRefSeq v211 references^101^ via vConTACT2 v2.0.11.3^41^ to infer approximately genus-level taxonomy of vOTUs (clustering together with references from a single genus); 2. manual inspection of the vConTACT2 clustering network (visualised in Cytoscape v3.8.2^102^) to infer relatedness at higher ranks (e.g. family or class) based on interactions with references in the protein sharing network; and 3. vConTACT2 clustering analysis of vOTUs together with high-quality sequences from the global IMG/VR database v7.1^84^. For all vConTACT2 steps, taxonomy assignments for viralRefSeq v211 references were reconciled with recent taxonomy updates (viralRefSeq v223)^101^ with higher ranks propagated based on ICTV taxonomy (MSL38.v3) (https://ictv.global/taxonomy; accessed 16 Sept 2023). To facilitate vConTACT2 analyses on the large dataset of IMG/VR references, sequences were first partitioned 50 ways, followed by 50 parallel vConTACT2 analyses together with vOTUs and viralRefSeq references (v211). For each partition, all IMG/VR sequences that shared an interaction in the resultant protein-sharing network were retained and concatenated into a final IMG/VR subset (*n*=131,283 sequences) and re-run through vConTACT2 to assess protein-sharing interactions with Waiwera estuary vOTUs. For the purposes of visualisation in Cytoscape v3.8.2^102^, the protein-sharing network was filtered to retain only IMG/VR sequences that shared an interaction with a vOTU (this study) or viralRefSeq sequence.

#### Inference of Caudoviricetes phylogeny via concatenated protein alignments of putative single copy core genes

Phylogeny of *Caudoviricetes* viruses (vOTUs, viralRefSeq, and IMG/VR sequences) was inferred via generating trees of concatenated filtered protein alignments of putative single copy “core genes” via Clustal Omega v1.2.4^103^, broadly following methodology previously described^53^, with core genes re-identified based on *Caudoviricetes* reference sequences in the viralRefSeq database v223^101^ (further details in **Supplementary Materials**). Phylogenetic trees based on the concatenated alignment were generated via IQ-TREE v2.2.2.2^104^ using ModelFinder (with VT+F+I+G4 selected)^105^, and with ultrafast bootstrap (nmax=1000 replicates)^106^. Trees were visualised using iTOL^107^. An alternative set of putative core genes was also identified for all high quality *Caudoviricetes* sequences (based on the provided taxonomy) in the IMG/VR database (**Supplementary Materials; Table S6**).

#### Identification of putative prokaryote hosts

Putative prokaryote hosts for Waiwera estuary vOTUs were predicted using a combination of approaches, including CRISPR-spacer, tRNA, and genome homology via pairwise blast searches (BLAST v2.13.0)^108^, analysis of vOTUs co-binned with prokaryote MAGs, oligonucleotide frequency similarity via VirHostMatcher v1.0.0^109^, and machine learning-based methods RaFAH v0.3^110^ and HostG (accessed 06 Dec 2021)^111^. These six methods were applied to the non-dereplicated MAG dataset and to all viral contigs, which were later summarised by vOTU cluster. Further details are available in **Supplementary Materials**.

#### vOTU and prokaryote ecology: spatial distribution, diversity, and ecological niche partitioning

vOTU and prokaryote MAG genome coverage (per million reads) and transcript abundances (per million reads) for each sample were calculated after mapping trimmed DNA and RNA reads, respectively, against all MAGs (n=646) and vOTUs (≥3 kb long, n=31,711) with BBMap v39.01^91^ (ambiguous=best minid=0.95), and filtered at a threshold of read counts <10 (genome) or <5 (transcriptome) (**Table S10; Table S11**). Genome coverage per million reads was calculated for each sample as follows: ((total mapped reads / genome length) / sequencing library size) * 1,000,000. RNA reads were assigned to genes via featureCounts (subread v2.0.6)^112^, and transcripts per million mapped reads (TPM) calculated via the following formula: (reads per-contig per-kilobase (RPK) / (sum(RPK) / 1,000,000)). MAGs and vOTUs were categorised into the following six distinct ecosystem types based on the sample in which the genome coverage was greatest (sites 3 and 7 were excluded to minimise edge effects associated with viral transport and mixing within the estuary): non-saline water or sediment (sites 1-2; 0.2-0.3 ppt); brackish water or sediment (sites 3-7; 20.2-26.3 ppt); and marine water or sediment (sites 8-9; 27.6-34.9 ppt).

Genome and transcriptome coverage heatmaps were generated in R v4.4.0^113^ using the *gplots* package^114^, with clustering dendrograms based on hierarchical clustering of weighted Bray-Curtis dissimilarity (calculated using the *vegan* package^115^). Alpha diversity (richness, Shannon [base=natural logarithm], and Simpson indices) of vOTUs and MAGs per assembly was also calculated using the *vegan* package. Beta diversity was assessed via non-metric multidimensional scaling (nMDS) ordinations and statistics (betadisper, TukeyHSD, and adonis2) based on weighted and unweighted Bray-Curtis dissimilarity in R using the *vegan* and *ggplot2*^116^ packages, with additional fitting of environmental variables. Changes in the phylogenetic relationships of prokaryote communities across the estuary were analysed via UniFrac analysis of concatenated alignments of prokaryote core genes (derived from GTDB-Tk) in R using the *phyloseq*^117^ and *ape*^118^ packages. Correlations between changes in the measured environmental variables, vOTU, and MAG community compositions, were assessed via Mantel tests comparing Bray-Curtis dissimilarites (vOTU and MAG communities) and Euclidean distances (environmental variables) using the *vegan* package in R.

The relatedness of virus genomes based on the environment types they were sampled from was assessed for Waiwera estuary vOTUs and high-quality environmental IMG/VR virus sequences (*n*=288,511) via two approaches: construction of a protein-sharing network and clustering with vConTACT2, and inferred phylogeny of *Caudoviricetes* viruses via alignment of putative core genes. Proportions of associated environment types for each vConTACT2-generated cluster were analysed and visualised in R. Trees of inferred *Caudoviricetes* phylogeny were generated as described above and manually inspected for associations between phylogenetic clustering and ecosystem types.

#### Gene GC and protein amino acid proportions and isoelectric point

Predicted genes and proteins from all MAGs, Waiwera vOTUs, and IMG/VR reference viruses were analysed to assess differential patterns in gene GC content and protein amino acid proportions and isoelectric points (pI) based on ecosystem type. Gene GC proportions were extracted from prodigal^119^ and prodigal-gv^120^ outputs. Amino acid proportions were calculated based on counts of individual amino acids within predicted protein sequences. Acidic, basic, polar, nonpolar, and charged amino acids were defined as per the program pepstats (EMBOSS v6.6.0)^121^. Delta acidic vs basic for each protein was calculated as proportion acidic – proportion basic amino acids. Isoelectric points for all predicted proteins were also calculated with pepstats. Genes and predicted proteins were categorised as major tail, tail tape measure, major capsid, endolysin, large terminase, DNA polymerase, spike, or ‘other’ by string matches and manual curation based on all annotations from DRAM-v for Waiwera vOTUs, and annotations based on viral orthologous groups database (VOGdb) (v222) matches via hmmsearch (HMMR v3.3.2; hmmer.org) for IMG/VR viruses. For the IMG/VR analyses, references were categorised based on the provided taxonomy affiliations. Linear models of individual VOGs identified across the estuary were generated in R v4.4.0 using the *stats* package^113^. Dot plots of GC and amino acid proportions against salinity were generated in R with the *ggplot2* package^116^, with linear modelling calculated via the *stat_poly_line()* and *stat_poly_eq()* functions from the *ggpmisc* package^122^. Ridge plots, density plots of pI, and dot plots of acidic, basic, polar, nonpolar, and charged amino acid proportions were generated using *ggplot2*. Comparisons across groups for IMG/VR data were calculated via Kruskal-Wallis rank sum tests^123^, followed by pairwise testing via Dunn’s tests with Bonferroni correction^124^.

#### Protein structural prediction and surface electrostatic potential

To investigate the influence of amino acid biases on protein surface electrostatic potentials, a subset of viruses was selected from three habitat types (non-saline and marine [Waiwera vOTUs], and hypersaline [IMG/VR]) for 3D structural prediction and analysis of four proteins of interest: major tail, major capsid, large terminase subunit, and DNA polymerase. Target proteins were first identified via word searches of DRAM-v annotations, with further manual curation based on structural prediction results. Viruses containing putative annotations for all four target proteins were selected. Six replicate viruses from each of the three habitat and four protein types were selected for further analysis. Three-dimensional structures of protein complexes (major tail and major capsid) and individual proteins (large terminase and DNA polymerase) were predicted from protein sequences via Alphafold 3 server^125^ with the default settings. Proteins were visualised in UCSF ChimeraX^126^, with electrostatic potential calculations via the *coulombic* function with a standardised range (coulombic range −30,30). Two putative large terminase proteins were excluded during analyses due to poor structural predictions, resulting in 70 proteins in total.

#### Changes in host translational machinery vs viral protein amino acid proportions

To investigate associations between tRNA gene and transcript abundances and amino acid proportions in predicted virus proteins, tRNA genes in prokaryotes and viral contigs were identified via DRAM v1.3.5^96^ and DRAM-v v1.4.6^96^, respectively. Mapping of metagenomic and metatranscriptomic reads to prokaryote and viral genomes, and abundance calculations, were conducted as described above. Relative proportions of prokaryote tRNA genes and transcripts were calculated to compare against proportions of encoded amino acids in predicted virus proteins. Overall changes in prokaryote tRNA repertoires were assessed via non-metric multidimensional scaling (nMDS) ordinations based on weighted Bray-Curtis dissimilarity in R using the *vegan* and *ggplot2*^116^ packages. Gene and transcript abundance heatmaps were generated in R v4.4.0^113^ using the *gplots* package^114^, with clustering dendrograms based on hierarchical clustering of weighted Bray-Curtis dissimilarity (calculated using the *vegan* package^115^). Dot and violin plots of tRNA gene and transcript proportions against salinity, and of virus contig length against virus tRNA counts per contig, were generated in R with the *ggplot2* package^116^, with linear modelling calculated via the *stat_poly_line()* and *stat_poly_eq()* functions from the *ggpmisc* package^122^. Spearman’s correlations comparing predicted virus protein amino acid proportions (for protein subsets and all proteins combined) against gene or transcript proportions of cognate prokaryote tRNAs were calculated using the *stats* package in R and results plotted via *ggplot2*.

#### Genome comparisons of selected closely related vOTUs

All-versus-all genome and protein alignments were generated with minimap2 v2.28^127^ (-X -N 50 -p 0.1) and blastp (BLAST v2.13.0)^108^ (-evalue 1e-5), respectively. Gene annotations generated by DRAM-v^96^ were categorised as auxiliary metabolic genes (AMG), replication- or structural-related (VOGdb categories Xr and Xs), and tail-, capsid-, or portal-related (via case-insensitive word searches of “tail”, “capsid”, or “portal” within VOGdb annotations generated by DRAM-v). Gene synteny plot was generated in R using the *gggenomes* package^128^. Phylogenetic trees, coverage heatmaps, dot plots of gene GC content and acidic-basic amino acid proportions, and density plots were generated and visualised as described above. Comparisons across all groups were calculated via Kruskal-Wallis rank sum tests^123^, followed by pairwise testing via Dunn’s tests with Bonferroni correction^124^.

## ACKNOWLEDGEMENTS

We thank G. Lear for contributions to the sampling plan, D. Waite, C. Astudillo-Garcia, G. Lear, J.S. Boey, and O.E. Mosley for assisting in sample collection, D. Waite for assistance with sample preparation and data processing, and I. Hay for guidance with protein structural prediction and analyses (University of Auckland). High-performance computing facilities were provided by New Zealand eScience Infrastructure.

## Authors’ Contributions

KH and MH conceived the study. HST, EG, and KH collected the samples, and HST generated the metagenomics assembly data. MH analysed the data with input from EG, JLG, and KH. MH and KH wrote the manuscript. All authors read and approved the final version of the manuscript.

## Data availability

Sequence data were deposited under NCBI BioProject PRJNA668816.

## Code availability

Workflow overview and custom scripts are available online at https://github.com/GenomicsAotearoa/workflow.DNA_virus_niche_adaptations.

## Ethics approval and consent to participate

Not applicable.

## SUPPLEMENTARY FIGURES

**Figure S1.** Rank abundance plots of within-vOTU microdiversity for (a) all vOTUs and (b, inset) the 50 most diverse vOTUs. Bars are coloured by the estuary zone that contigs within vOTUs were derived from. (c, inset) Beeswarm plots of within-vOTU microdiversity, with vOTUs categorised by the estuary zone that the vOTU was most abundant in. (d) Dot plots of alpha diversity indices (richness, Shannon (base = natural logarithm), Simpson, and inverse Simpson) against sample salinity for vOTUs (top row; *n*=858) and prokaryote MAGs (bottom row; *n*=646). Lines (with 95% confidence interval shaded) calculated via loess local smoothing.

**Figure S2.** vOTU and prokaryote MAG distributions throughout the Waiwera estuary. Non-metric Multidimensional Scaling (nMDS) ordinations comparing the composition of vOTUs (>50% complete, left column [*n*=858]; all vOTUs, centre column [*n*=31,711]) and MAGs (rightmost plots, *n*=646) per sample. The ordinations are based on weighted and unweighted Bray-Curtis dissimilarities using (a) whole genome sequencing data (genome relative abundance) and (b) transcriptome sequencing data. Arrows represent fitting of environmental parameters.

**Figure S3.** Viral clusters and phylogeny by habitat. (a) Clusters generated by vConTACT2 containing viruses from more than one ecosystem type (non-saline, saline, or terrestrial). Counts (top row) and percentages (middle and bottom rows) of viruses recovered from these ecosystems. Clusters contain viruses from IMG/VR (all rows), and at least one Waiwera vOTU (top and middle rows). Cluster counts, percentage of total clusters (parentheses), and virus sequence counts per group: Non-saline+Terrestrial clusters=14 (5.0%), sequences=214; Non-saline+Saline+Terrestrial clusters=30 (10.7%), sequences=1,001; Non-saline+Saline clusters=65 (23.2%), sequences=1,895; Saline+Terrestrial clusters=8 (2.9%), sequences=55. All mixed clusters (including Waiwera vOTUs and high-quality IMG/VR sequences) are presented in the bottom row. Cluster counts (and percentage of total clusters) and virus sequence counts per group: Non-saline+Terrestrial clusters=1,379 (5.6%), sequences=14,945; Non-saline+Saline+Terrestrial clusters=713 (2.9%), sequences=13,759; Non-saline+Saline clusters=1,746 (7.1%), sequences=24,692; Saline+Terrestrial clusters=504 (2.0%), sequences=4,834. (b) Inferred *Caudoviricetes* phylogeny for vOTUs (this study) and viralRefSeq references based on TerL protein alignment. Lines extending from branch tips for Waiwera vOTUs are coloured by estuary zone. Estuary sample type represents the estuary zone where the vOTU was most abundant. The outer ring is coloured by predicted vOTU and RefSeq taxonomy. (c) Inferred *Caudoviricetes* phylogeny for high-quality environmental viruses from the IMG/VR database based on concatenated alignment of predicted core genes. Putative core genes were identified using high-quality IMG/VR *Caudoviricetes* viruses as the initial reference set. The number of concatenated proteins used was 13. Lines extending from branch tips are coloured by habitat. (b,c) Trees were generated via IQ-TREE using ModelFinder (with VT+F+I+G4 selected). The scale bars represent amino acid substitutions per site.

**Figure S4.** Molecular signatures of environmental adaptation in DNA viruses from the Waiwera estuary. (a) Proportions of acidic, basic, polar, nonpolar, and charged amino acids, and the difference between the proportion of acidic-basic amino acids, in predicted viral contig protein subsets and all combined proteins in the estuary water column. Linear modelling equation and statistics were calculated via the *stat_poly_line()* and *stat_poly_eq()* functions from the *ggpmisc* package in R. (b) Predicted protein isoelectric points against salinity for predicted viral contig protein subsets and all combined proteins in the estuary water column. (c) x-estimates from linear models showing the trend of changes in individual amino acid proportions with respect to increases in salinity across the estuary in water, with bars representing median and interquartile range. Plots represent orthologous groups per protein type with a minimum of 20 sequences/ortholog. Numbers of orthologous groups (and sequences) per protein type: major tail = 7 (364); tail tape measure = 4 (201); major capsid =10 (3,272); endolysin = 9 (2,131); TerL = 7 (2,874); DNA polymerase = 17 (4,958); and total = 1,230 (184,695). (d) Mean GC content of amino acid codons against x-estimates from linear models showing the trend of changes in amino acid proportions with respect to increases in salinity across the estuary in water. (e) x-estimates from linear models showing the trend of changes in viral contig gene GC content for individual orthologous groups (>20 sequences/ortholog) with respect to increases in salinity across the estuary in water, with bars representing median and interquartile range. VOGs with *p*-values <0.05 in individual linear models are shown.

**Figure S5.** Front and back views of three-dimensional structural predictions of viral protein complexes (major tail and major capsid) and individual proteins (large terminase and DNA polymerase) colored by surface electrostatic potentials (ESP; blue to red surface coloring) for six representatives each from non-saline, marine, and hypersaline environments. Proteins from non-saline and marine habitats were from the Waiwera estuary. Those from hypersaline habitats were from IMG/VR. Protein structures were predicted from protein sequences via Alphafold 3 server, with visualisation and ESP calculation in ChimeraX. ESP was calculated via the *coulombic* function with a standardised range (coulombic range −30,30). Two putative large terminase proteins were excluded during analyses due to poor structural predictions, resulting in 70 proteins in total.

**Figure S6.** Molecular signatures of environmental adaptation in DNA viruses from the IMG/VR database. Amino acid composition (a) and isoelectric points (pI) (b) of predicted proteins per ecosystem type split by functional category (a and b) and viral realm (a only) for high-quality IMG/VR viruses. Number of predicted genes per functional category per realm are shown on plots (n=).

**Figure S7.** Molecular signatures of environmental adaptation in DNA viruses from the IMG/VR database. (a) GC content per functional category per ecosystem type for high-quality IMG/VR viruses. (b) GC content per ecosystem type (summarised into broad ecosystem categories) split by functional category and viral realm for high-quality IMG/VR viruses.

**Figure S8**. Prokaryote tRNA gene counts. (a) Non-Metric Multidimensional Scaling ordination comparing samples based on Bray-Curtis dissimilarities calculated from prokaryote MAG tRNA gene abundances. (b-c) Counts of tRNA genes for each amino acid in all prokaryote MAGs (*n*=646) (b), and in Luna cluster prokaryote MAGs (*n*=19) (c). Inner colour bars represent taxonomy. Outer colour bars represent assigned estuary zone based on the site where the MAG was most abundant.

**Figure S9.** Proportions of prokaryote tRNA gene (a) and transcript (b) abundances in each MAG against salinity in the Waiwera estuary water column for all amino acids. Linear modelling equations and statistics were calculated via the *stat_poly_line()* and *stat_poly_eq()* functions from the *ggpmisc* package in R, with line shading representing 95% confidence interval. (c) Spearman’s correlations comparing predicted virus protein amino acid and cognate prokaryote tRNA gene (left panel) and transcript (right panel) relative proportions across the Waiwera estuary water samples for virus protein subsets and all proteins combined. * = Spearman’s *p*-value ≤0.05.

**Figure S10.** Molecular signatures of environmental adaptation in DNA viruses from the Waiwera estuary. (a) Proportions of glutamic acid, aspartic acid, arginine, histidine, lysine, and alanine in predicted viral contig protein subsets and all combined proteins in the estuary water column. Linear modelling equation and statistics were calculated via the *stat_poly_line()* and *stat_poly_eq()* functions from the *ggpmisc* package in R.

**Figure S11.** Traits of closely related vOTUs and host predictions. (a) Comparison of genes/proteins in common (>40% AA identity) for the two closely related vOTUs with complete genomes that collectively span the salinity divide. Comparisons are based on protein isoelectric points, the difference between encoding of acidic-basic amino acids, and gene GC content. Bars in dot plots represent the median and interquartile range. Each dot represents values for individual genes or proteins. (b-c) vOTU-prokaryote host (MAG) matches based on CRISPR spacer blast searches at 100% identity. (b) Counts and taxonomy (phylum) of MAG matches per vOTU. (c) Counts of vOTU matches per prokaryote MAG. Smaller plots are after filtering for MAGs with CRISPR regions further verified via CRISPRDetect.

## SUPPLEMENTARY TABLES

Table S1. Statistics of viral contigs identified for each assembly

Table S2. vOTU details of all vOTUs predicted to be ≥50% complete

Table S3. Total counts of vOTUs with taxonomy predicted

Table S4. Counts per taxonomic rank of vOTUs with taxonomy predicted

Table S5. Results of simple and partial Mantel tests

Table S6. Predicted *Caudoviricetes* single copy core gene sets

Table S7. Additional statistics results summaries

Table S8. Sequencing and assembly statistics

Table S9. Prokaryote metagenome-assembled genomes statistics

Table S10. Genome coverage per sample for MAGs and vOTUs

Table S11. vOTU gene transcripts per million per sample and gene annotations

Table S12. CRISPR spacer host matches for vOTUs

