## Supplementary Figures for "Adaptations of DNA viruses are influenced by host and environment with proliferations constrained by environmental niche"

Short title: DNA virus niche adaptations

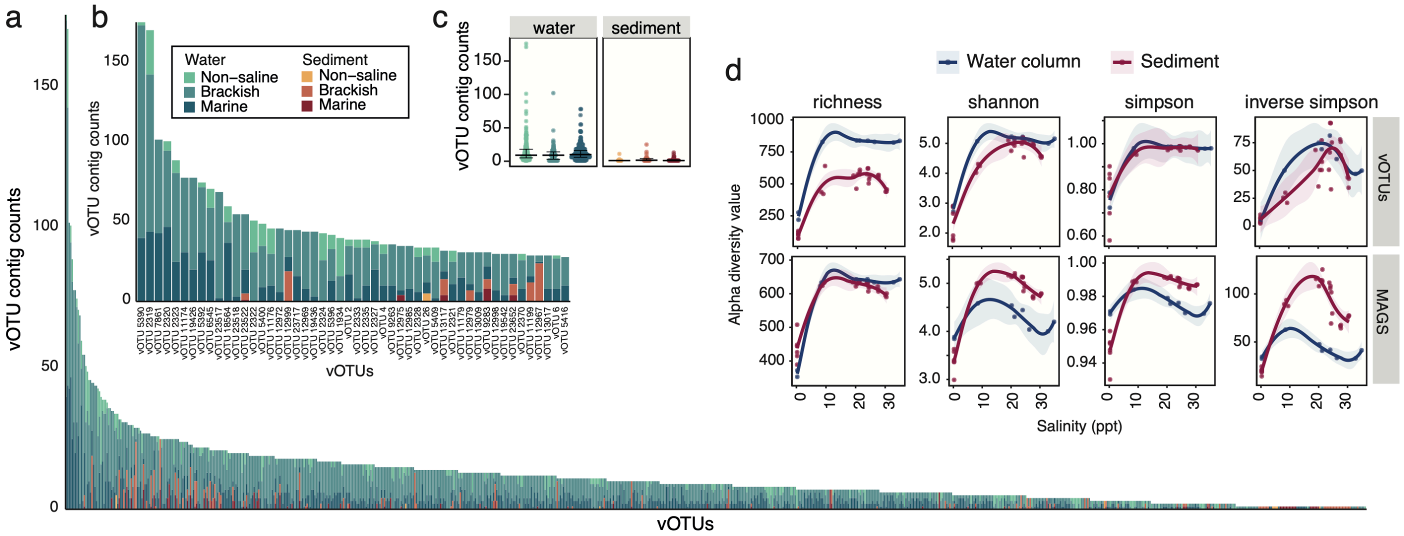


**Figure S1.** Rank abundance plots of within-vOTU microdiversity for (a) all vOTUs and (b, inset) the 50 most diverse vOTUs. Bars are coloured by the estuary zone that contigs within vOTUs were derived from. (c, inset) Beeswarm plots of within-vOTU microdiversity, with vOTUs categorised by the estuary zone that the vOTU was most abundant in. (d) Dot plots of alpha diversity indices (richness, Shannon (base = natural logarithm), Simpson, and inverse Simpson) against sample salinity for vOTUs (top row; $n$=858) and prokaryote MAGs (bottom row; $n$=646). Lines (with 95% confidence interval shaded) calculated via loess local smoothing.

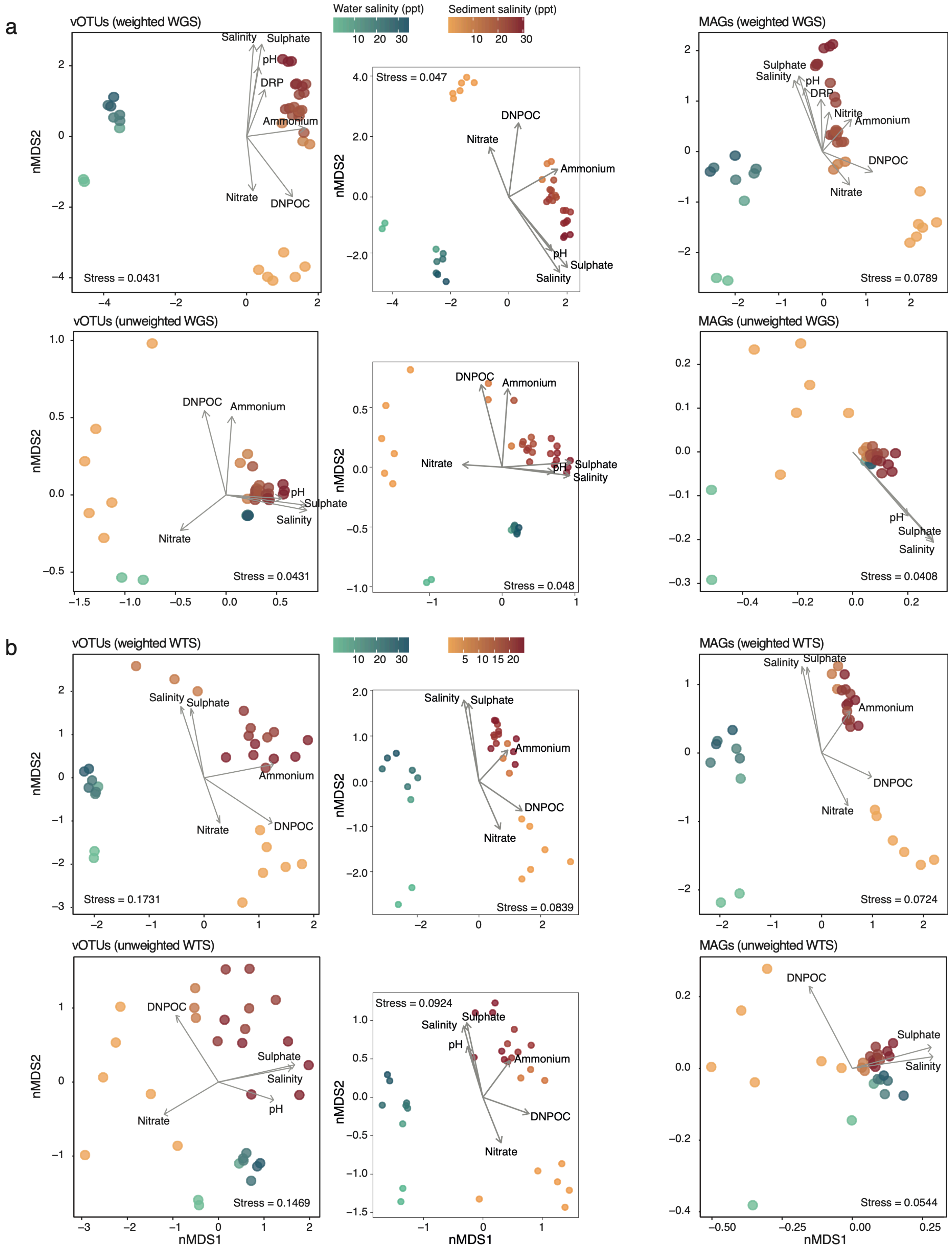


**Figure S2.** vOTU and prokaryote MAG distributions throughout the Waiwera estuary. Non-metric Multidimensional Scaling (nMDS) ordinations comparing the composition of vOTUs (>50% complete, left column [*n*=858]; all vOTUs, centre column [*n*=31,711]) and MAGs (rightmost plots, *n*=646) per sample. The ordinations are based on weighted and unweighted Bray-Curtis dissimilarities using (a) whole genome sequencing data (genome relative abundance) and (b) transcriptome sequencing data. Arrows represent fitting of environmental parameters.

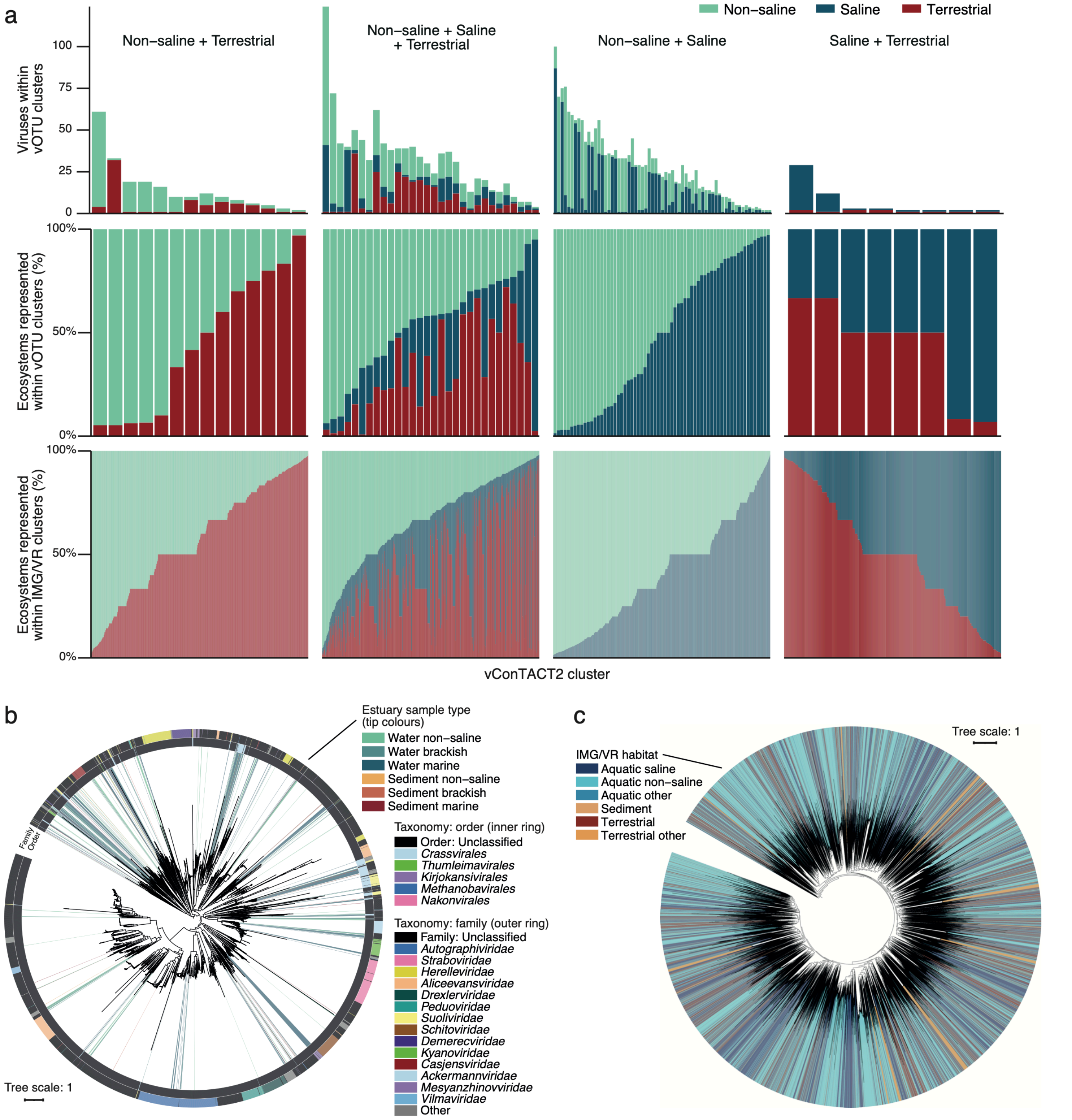


**Figure S3.** Viral clusters and phylogeny by habitat. (a) Clusters generated by vConTACT2 containing viruses from more than one ecosystem type (non-saline, saline, or terrestrial). Counts (top row) and percentages (middle and bottom rows) of viruses recovered from these ecosystems. Clusters contain viruses from IMG/VR (all rows), and at least one Waiwera vOTU (top and middle rows). Cluster counts, percentage of total clusters (parentheses), and virus sequence counts per group: Non-saline+Terrestrial clusters=14 (5.0%), sequences=214; Non-saline+Saline+Terrestrial clusters=30 (10.7%), sequences=1,001; Non-saline+Saline clusters=65 (23.2%), sequences=1,895; Saline+Terrestrial clusters=8 (2.9%), sequences=55. All mixed clusters (including Waiwera vOTUs and high-quality IMG/VR sequences) are presented in the bottom row. Cluster counts (and percentage of total clusters) and virus sequence counts per group: Non-saline+Terrestrial clusters=1,379 (5.6%), sequences=14,945; Non-saline+Saline+Terrestrial clusters=713 (2.9%), sequences=13,759; Non-saline+Saline clusters=1,746 (7.1%), sequences=24,692; Saline+Terrestrial clusters=504 (2.0%), sequences=4,834. (b) Inferred *Caudoviricetes* phylogeny for vOTUs (this study) and viralRefSeq references based on TerL protein alignment. Lines extending from branch tips for Waiwera vOTUs are coloured by estuary zone. Estuary sample type represents the estuary zone where the vOTU was most abundant. The outer ring is coloured by predicted vOTU and RefSeq taxonomy. (c) Inferred *Caudoviricetes* phylogeny for high-quality environmental viruses from the IMG/VR database based on concatenated alignment of predicted core genes. Putative core genes were identified using high-quality IMG/VR *Caudoviricetes* viruses as the initial reference set. The number of concatenated proteins used was 13. Lines extending from branch tips are coloured by habitat. (b,c) Trees were generated via IQ-TREE using ModelFinder (with VT+F+I+G4 selected). The scale bars represent amino acid substitutions per site.

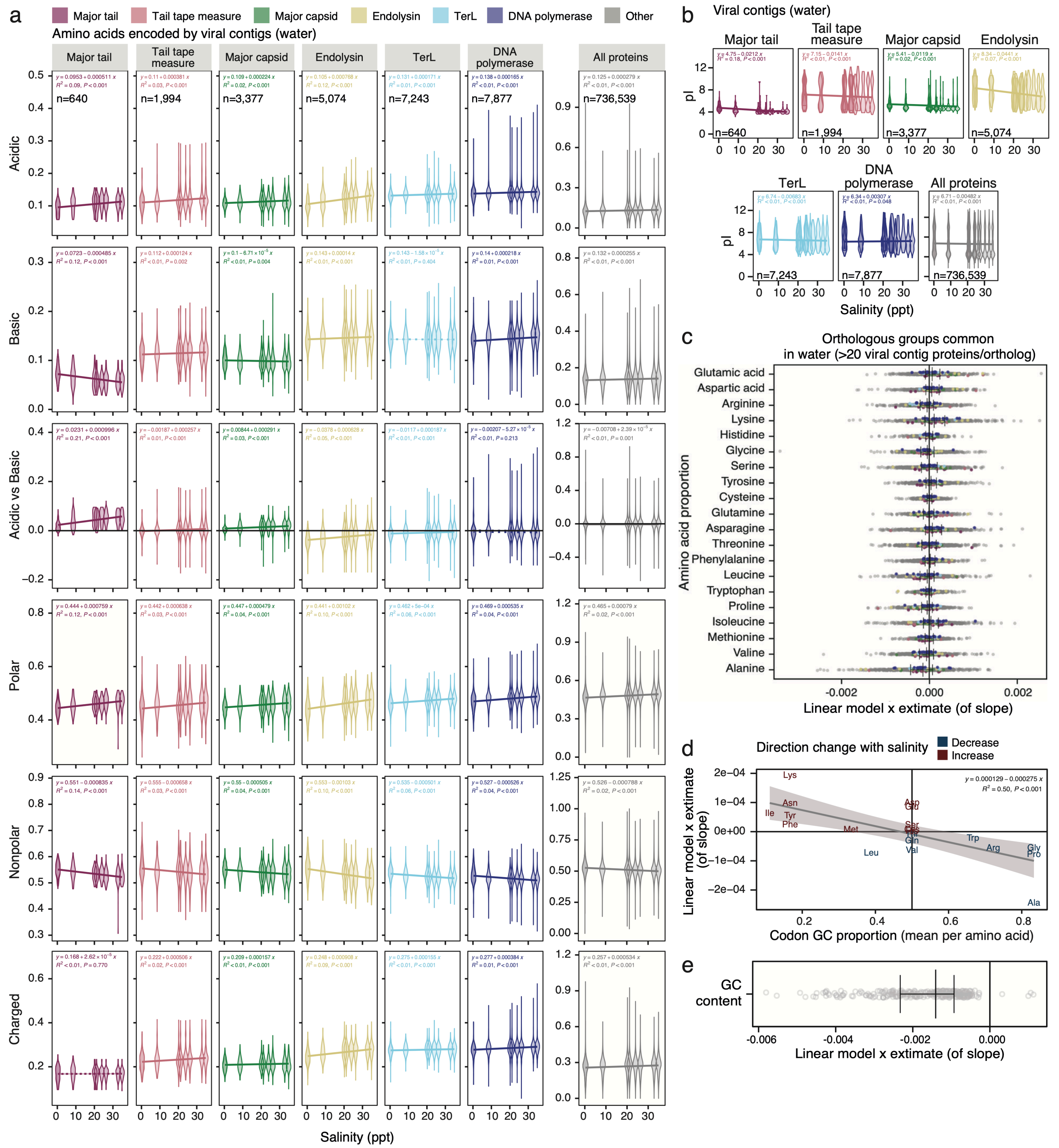


**Figure S4.** Molecular signatures of environmental adaptation in DNA viruses from the Waiwera estuary. (a) Proportions of acidic, basic, polar, nonpolar, and charged amino acids, and the difference between the proportion of acidic-basic amino acids, in predicted viral contig protein subsets and all combined proteins in the estuary water column. Linear modelling equation and statistics were calculated via the *stat_poly_line()* and *stat_poly_eq()* functions from the *ggpmisc* package in R. (b) Predicted protein isoelectric points against salinity for predicted viral contig protein subsets and all combined proteins in the estuary water column. (c) x-estimates from linear models showing the trend of changes in individual amino acid proportions with respect to increases in salinity across the estuary in water, with bars representing median and interquartile range. Plots represent orthologous groups per protein type with a minimum of 20 sequences/ortholog. Numbers of orthologous groups (and sequences) per protein type: major tail = 7 (364); tail tape measure = 4 (201); major capsid =10 (3,272); endolysin = 9 (2,131); TerL = 7 (2,874); DNA polymerase = 17 (4,958); and total = 1,230 (184,695). (d) Mean GC content of amino acid codons against x-estimates from linear models showing the trend of changes in amino acid proportions with respect to increases in salinity across the estuary in water. (e) x-estimates from linear models showing the trend of changes in viral contig gene GC content for individual orthologous groups (>20 sequences/ortholog) with respect to increases in salinity across the estuary in water, with bars representing median and interquartile range. VOGs with p-values <0.05 in individual linear models are shown.

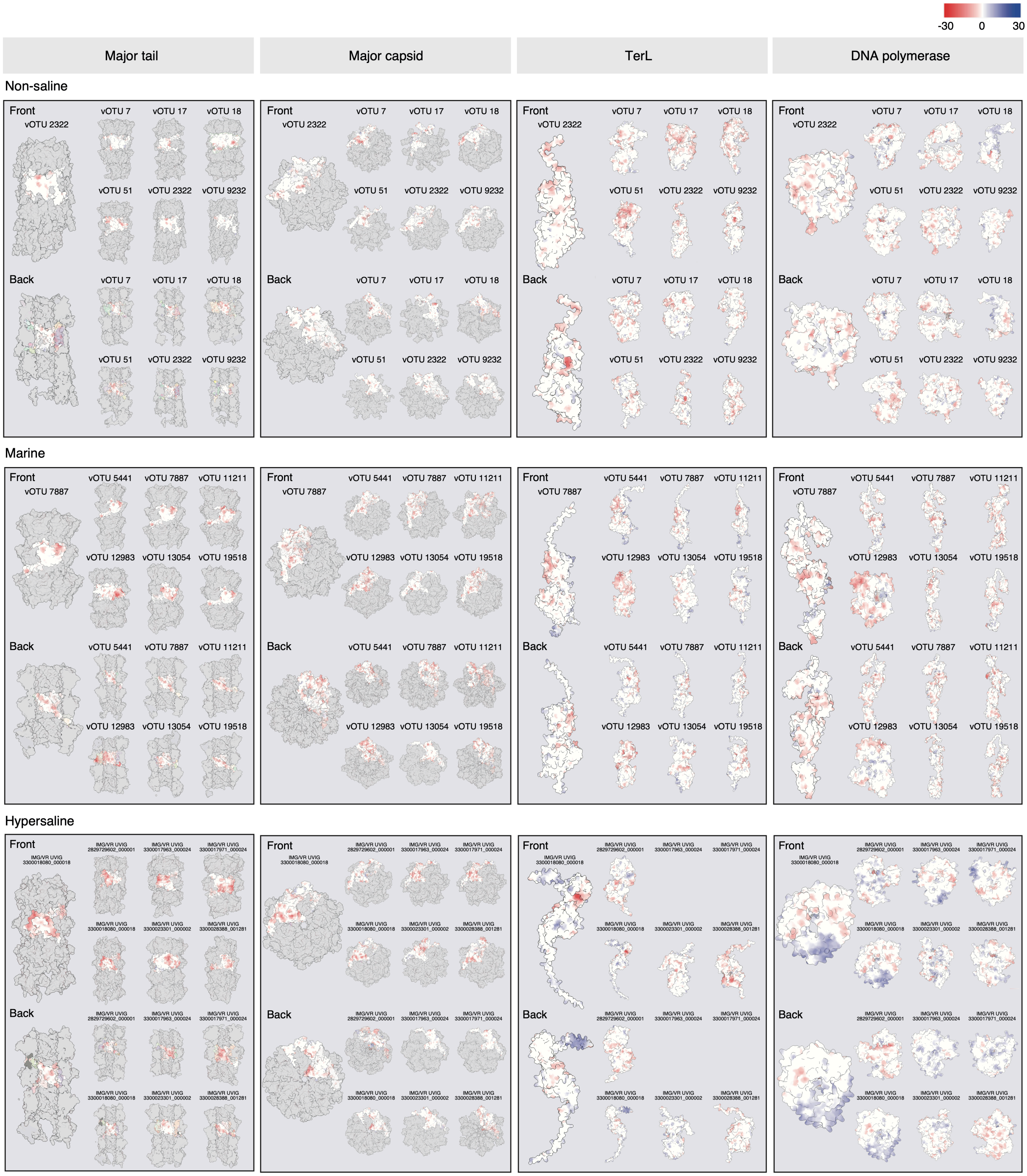


**Figure S5.** Front and back views of three-dimensional structural predictions of viral protein complexes (major tail and major capsid) and individual proteins (large terminase and DNA polymerase) colored by surface electrostatic potentials (ESP; blue to red surface coloring) for six representatives each from non-saline, marine, and hypersaline environments. Proteins from non-saline and marine habitats were from the Waiwera estuary. Those from hypersaline habitats were from IMG/VR. Protein structures were predicted from protein sequences via Alphafold 3 server, with visualisation and ESP calculation in ChimeraX. ESP was calculated via the *coulombic* function with a standardised range (coulombic range -30,30). Two putative large terminase proteins were excluded during analyses due to poor structural predictions, resulting in 70 proteins in total.

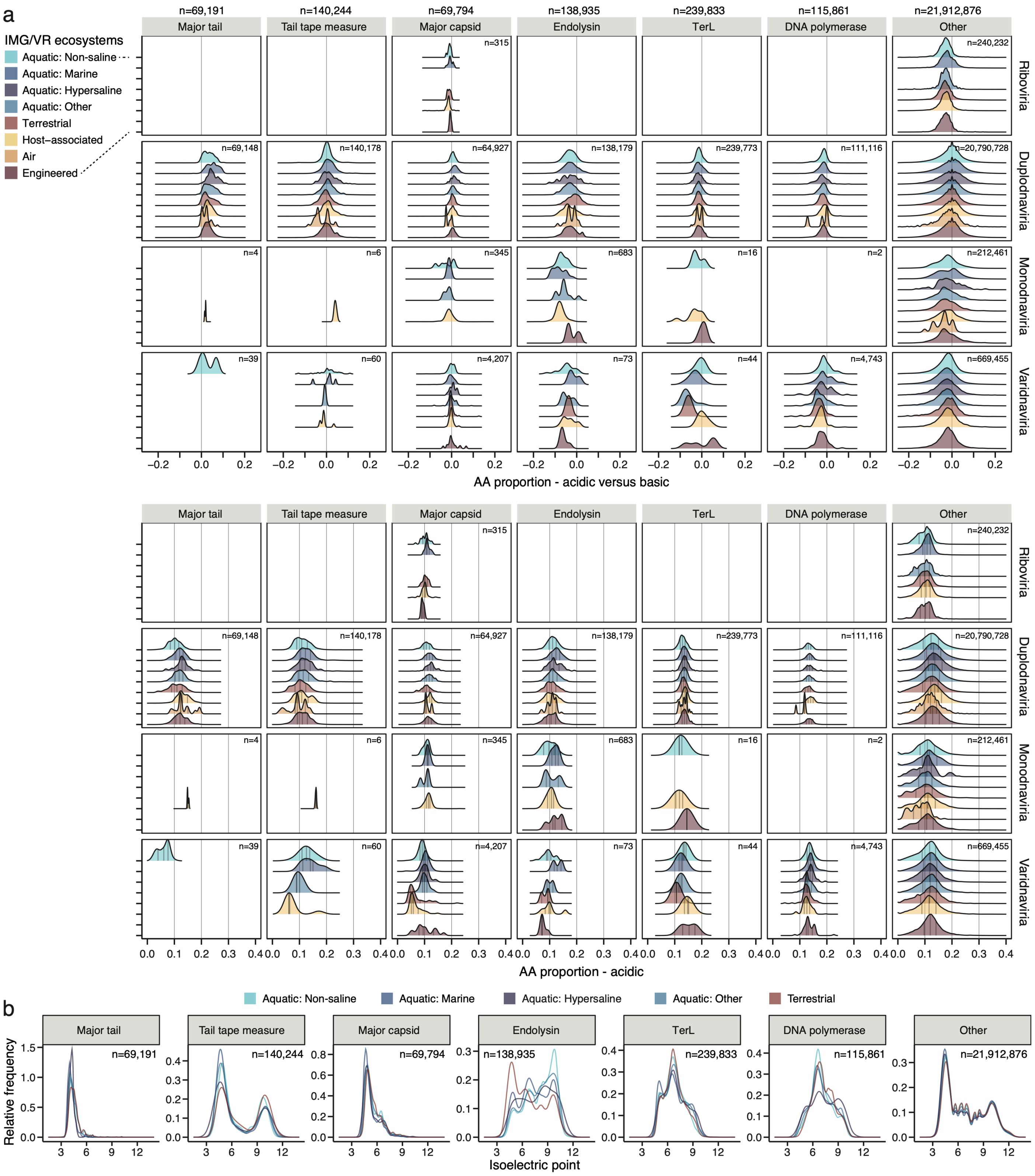


**Figure S6.** Molecular signatures of environmental adaptation in DNA viruses from the IMG/VR database. Amino acid composition (a) and isoelectric points (pI) (b) of predicted proteins per ecosystem type split by functional category (a and b) and viral realm (a only) for high-quality IMG/VR viruses. Number of predicted genes per functional category per realm are shown on plots (n=).

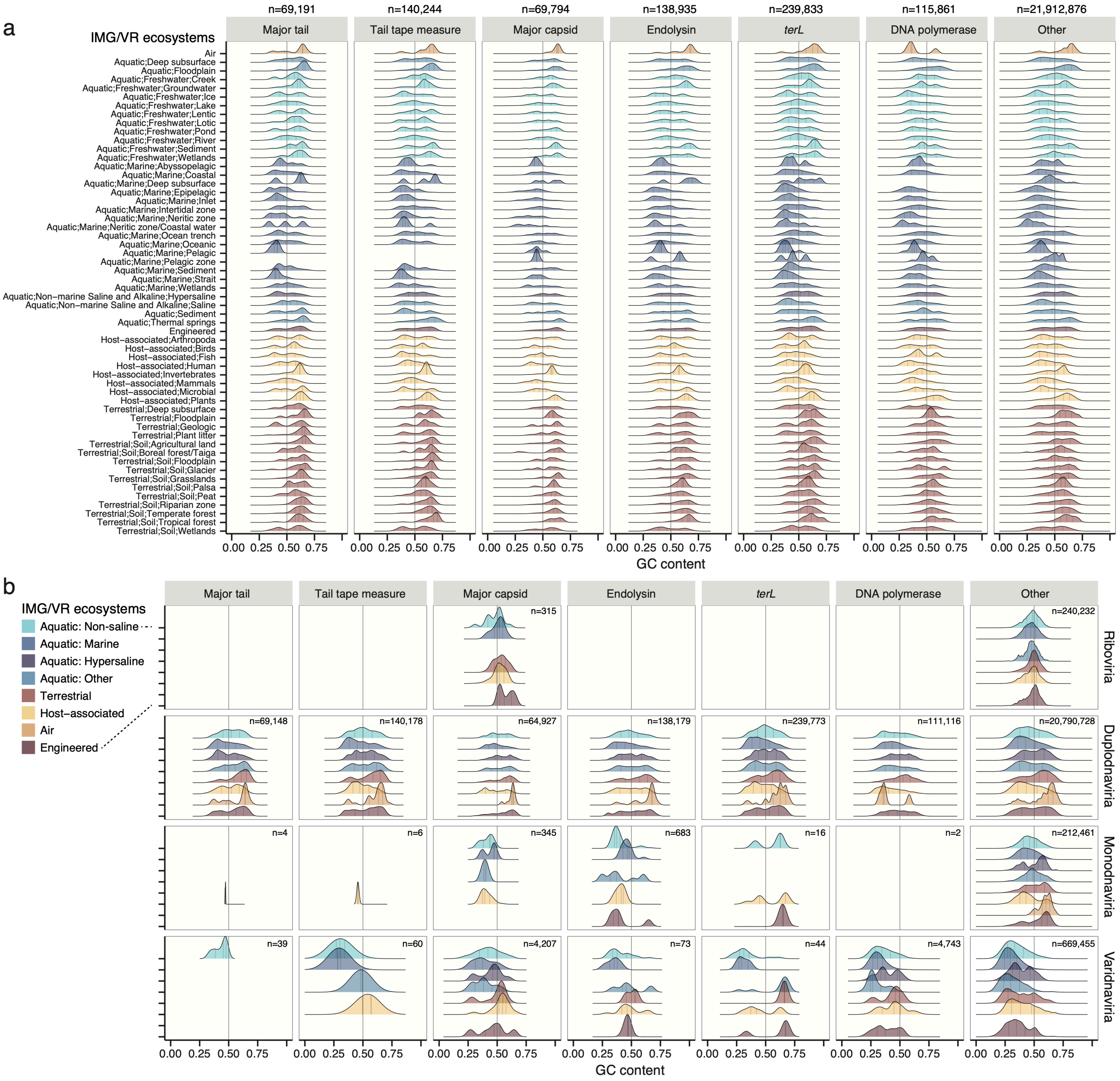


**Figure S7.** Molecular signatures of environmental adaptation in DNA viruses from the IMG/VR database. (a) GC content per functional category per ecosystem type for high-quality IMG/VR viruses. (b) GC content per ecosystem type (summarised into broad ecosystem categories) split by functional category and viral realm for high-quality IMG/VR viruses.

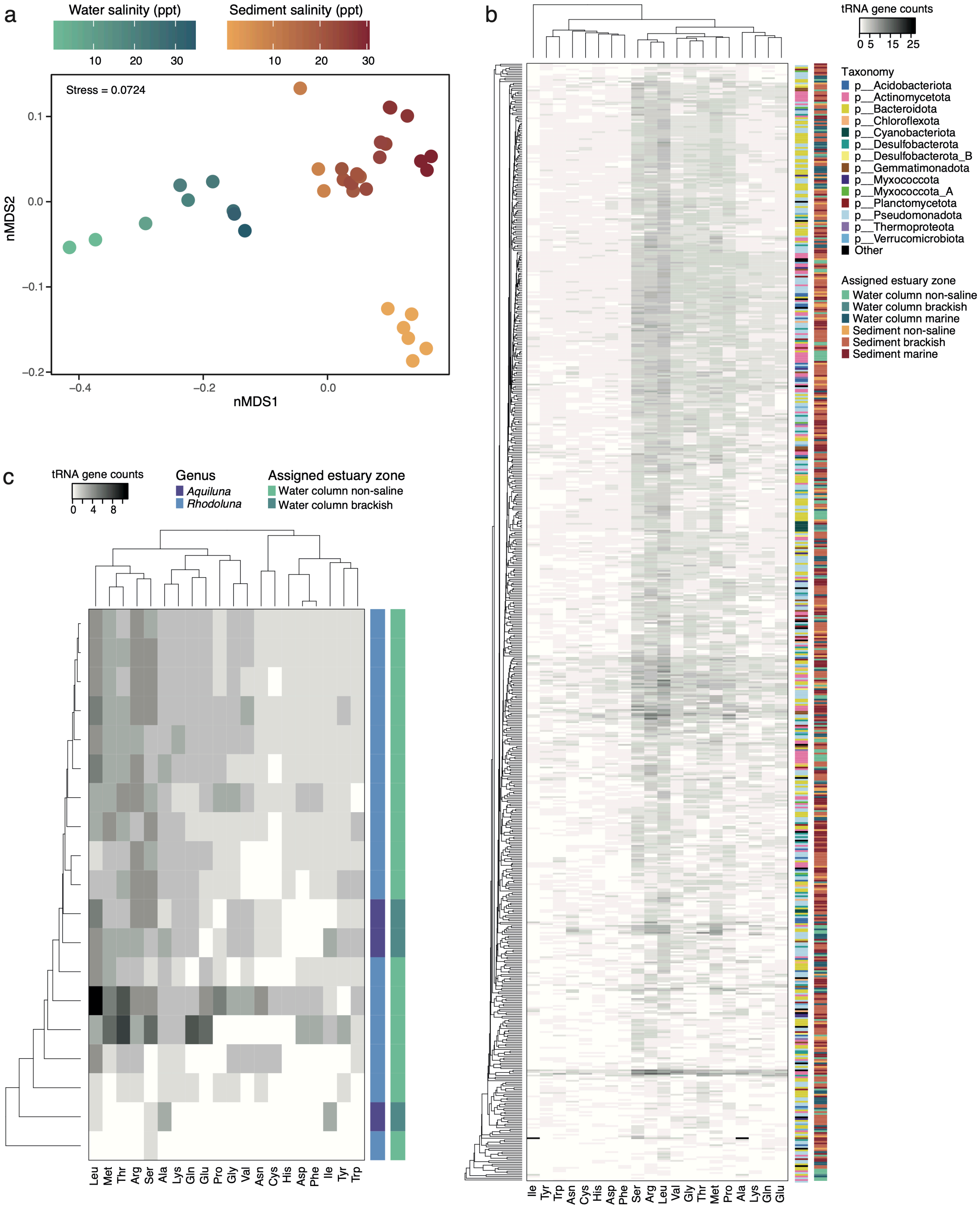


**Figure S8.** Prokaryote tRNA gene counts. (a) Non-Metric Multidimensional Scaling ordination comparing samples based on Bray-Curtis dissimilarities calculated from prokaryote MAG tRNA gene abundances. (b-c) Counts of tRNA genes for each amino acid in all prokaryote MAGs (*n*=646) (b), and in Luna cluster prokaryote MAGs (*n*=19) (c). Inner colour bars represent taxonomy. Outer colour bars represent assigned estuary zone based on the site where the MAG was most abundant.

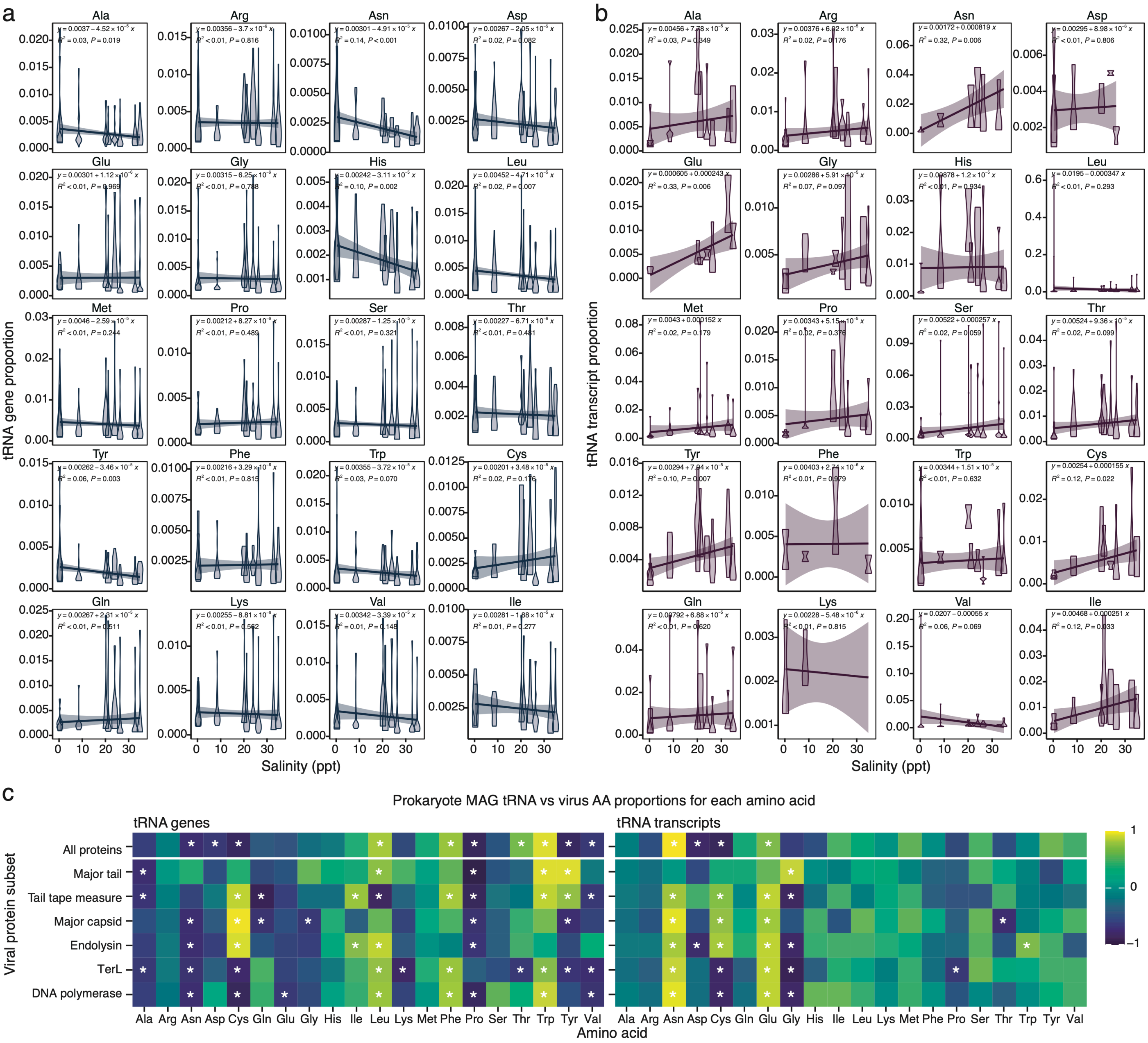


**Figure S9.** Proportions of prokaryote tRNA gene (a) and transcript (b) abundances in each MAG against salinity in the Waiwera estuary water column for all amino acids. Linear modelling equations and statistics were calculated via the *stat_poly_line()* and *stat_poly_eq()* functions from the *ggpmisc* package in R, with line shading representing 95% confidence interval. (c) Spearman's correlations comparing predicted virus protein amino acid and cognate prokaryote tRNA gene (left panel) and transcript (right panel) relative proportions across the Waiwera estuary water samples for virus protein subsets and all proteins combined. * = Spearman's *p*-value ≤0.05.

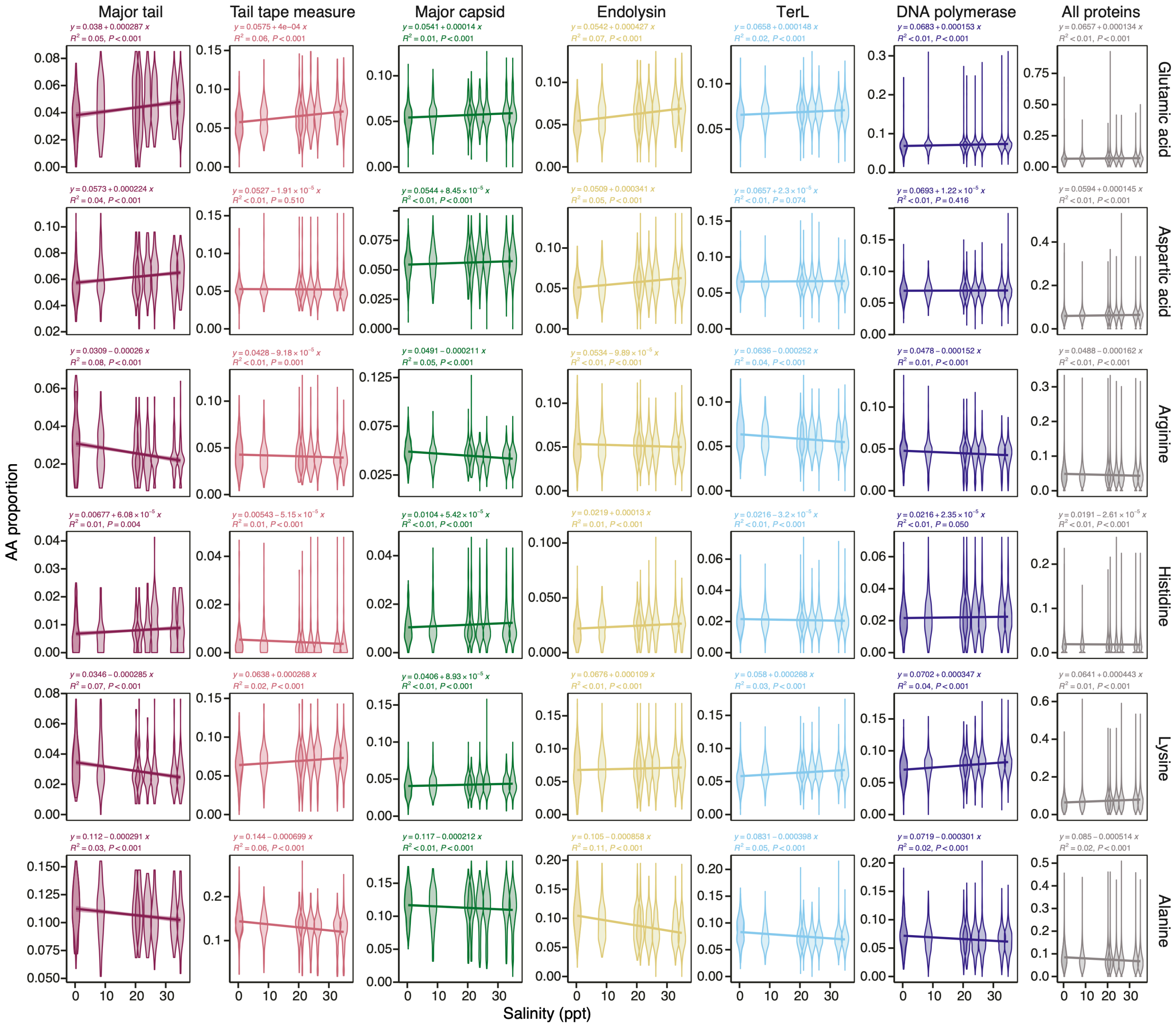


**Figure S10.** Molecular signatures of environmental adaptation in DNA viruses from the Waiwera estuary. (a) Proportions of glutamic acid, aspartic acid, arginine, histidine, lysine, and alanine in predicted viral contig protein subsets and all combined proteins in the estuary water column. Linear modelling equation and statistics were calculated via the *stat_poly_line()* and *stat_poly_eq()* functions from the *ggpmisc* package in R.

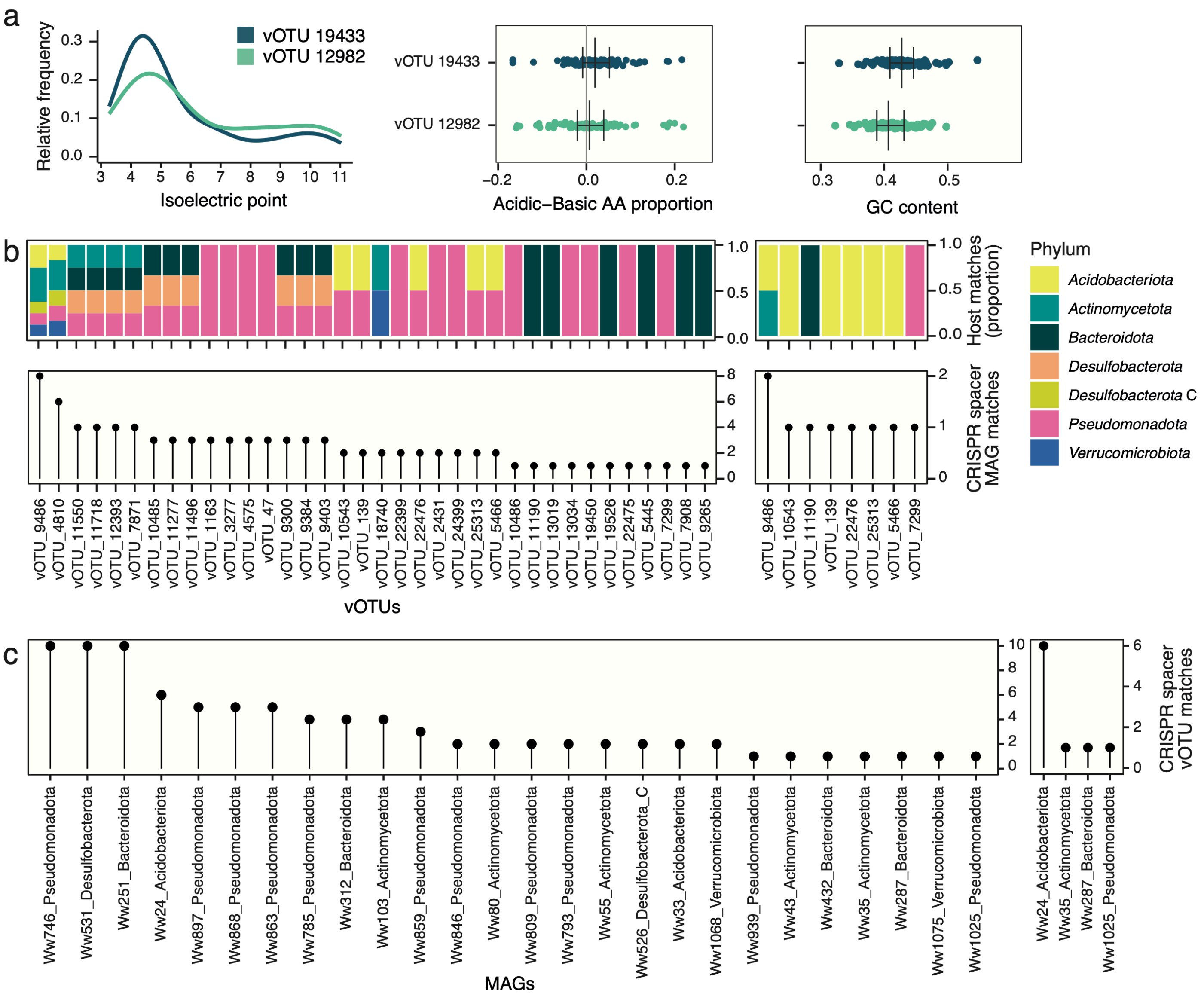


**Figure S11.** Traits of closely related vOTUs and host predictions. (a) Comparison of genes/proteins in common (>40% AA identity) for the two closely related vOTUs with complete genomes that collectively span the salinity divide. Comparisons are based on protein isoelectric points, the difference between encoding of acidic-basic amino acids, and gene GC content. Bars in dot plots represent the median and interquartile range. Each dot represents values for individual genes or proteins. (b-c) vOTU-prokaryote host (MAG) matches based on CRISPR spacer blast searches at 100% identity. (b) Counts and taxonomy (phylum) of MAG matches per vOTU. (c) Counts of vOTU matches per prokaryote MAG. Smaller plots are after filtering for MAGs with CRISPR regions further verified via CRISPRDetect.
