## Supplementary Materials for "Adaptations of DNA viruses are influenced by host and environment with proliferations constrained by environmental niche"

Short title: DNA virus niche adaptations

#### SUPPLEMENTARY RESULTS

##### Host matching remains a significant challenge in viral metagenomics studies

We assessed the putative prokaryote host range of the vOTUs in our estuary data set using a combination of approaches, including CRISPR-spacer, tRNA, and genome pairwise blast searches, analysis of vOTUs co-binned with prokaryote MAGs, VirHostMatcher, and machine learning-based methods RaFAH and HostG. Inconsistent results, including up to the rank of phylum, were frequently observed among the different methods. Previous studies have applied a ranking system across multiple methods based on the reliability of the method, with, for example, CRISPR-spacer matches out-ranking matches based on tRNA or genome blast searches[1,2]. However, the inconsistencies observed here for predicted viral host matches question how robust inferences are based on many of the currently applied approaches. As CRISPR-spacer matches are considered the most robust method in metagenomics-based phage host-matching[2], our remaining analyses were restricted to this subset of results. Based on CRISPR spacer analysis, 121 Waiwera vOTUs (4 vOTUs ≥50% complete) were putatively matched to 28 prokaryote MAGs in this dataset, with viral sequences within individual vOTUs matching between 1 and 11 (mean = 3.1) prokaryote MAGs and individual MAGs matching between 1 and 90 vOTUs (**Figure S11b,c; Table S12**). CRISPR spacer matches for viral sequences within individual vOTUs also often spanned bacterial phyla, with *Pseudomonadota* returning the most frequent matches. Further filtering to retain putative hosts where CRISPR arrays were also identified using CRISPRDetect resulted in matches remaining for 8 vOTUs (1 vOTU ≥50% complete) to 4 MAGs (**Figure S11b,c**).

### SUPPLEMENTARY METHODS

#### Data acquisition

*Waiwera estuary metagenomics and metatranscriptomics*

Metagenomic and metatranscriptomic Illumina sequencing was conducted on filtered water and triplicate sediment samples from nine sites spanning the salinity gradient and large differences in nutrient availability (water n = 9 samples; sediment n = 9 x 3 = 27 samples), as described previously[3]. In brief, filtered water and triplicate sediment samples were collected in November 2018 during low tide from nine sites spanning the salinity gradient (0.2 to 35 ppt) of the Waiwera river and estuary, Aotearoa New Zealand (36° 33′ 02.4″ S, 174° 39′ 08.1″ E). Water samples (~10L) were sequentially filtered through sterile 1.2 µm and 0.22 µm mixed cellulose ester filters (Merck Millipore, MA, USA). The viral fraction of these samples includes those associated with cells or biofilms on particles trapped on 0.22 micron filters. Triplicate wet-sieved (1 mm mesh sieve) sediment samples ~5 m apart were collected to a depth of 2 cm at each of the nine sites. Samples were placed in sterile 50 mL Falcon tubes, preserved in LifeGuard Soil Preservation Solution (Qiagen, MD, USA), and stored at -80 °C. Additional water and sediment samples were also collected for analyses of physicochemical and nutrient composition as previously described[3].

Illumina HiSeq (2500) paired end 2x250 bp and 2x125 bp sequencing of extracted total DNA and RNA (RNeasy PowerSoil Total RNA and RNeasy PowerSoil DNA Elution kits (Qiagen, MD, USA)), respectively, was conducted by the Otago Genomics Facility (University of Otago, Dunedin, New Zealand). Metagenomics reads were quality trimmed and filtered with Trimmomatic (v0.38)[4] and assembled with metaSPAdes (v3.11.1)[5] (**Table S8**). Individual assemblies were generated for each sample type (water and sediment) and site (*n* = 9) from the estuary (triplicate sediment samples for each site were pooled prior to assembly), generating 18 metagenome assemblies.

#### Data processing

*Inference of Caudoviricetes phylogeny via concatenated protein alignments of putative single copy core genes*

Phylogeny of *Caudoviricetes* viruses (vOTUs, viralRefSeq, and IMG/VR sequences) was inferred via generating trees of concatenated protein alignments of putative single copy "core genes" broadly following methodology previously described[6]. The VOG database has expanded since this previous work, and marker gene IDs identified there are no longer applicable. As such, the process of selecting core genes and generating a concatenated gene tree of alignments was reproduced here, with core genes re-identified based on *Caudoviricetes* reference sequences in the viralRefSeq database (**Table S6**). Genomic sequences were downloaded from the viralRefSeq database v223[7], filtered to retain putative Caudoviricetes sequences based on the provided sequence taxonomy and sequences with predicted genome completeness ≥ 95% (based on CheckV analysis[8]), and genes predicted and annotated via DRAM-v v1.4.6[9]. Predicted proteins for all viralRefSeq reference genomes and vOTUs from this study were searched via hmmsearch (HMMR v3.3.2; hmmer.org) against profile hidden Markov models (HMMs) for viral proteins downloaded from the viral orthologous groups database v222 (VOGdb) with a threshold of -E 1e-3. For each geneID, VOGdb matches with the lowest full sequence evalue score were retained. Putative single copy core genes were identified based on the following criteria: 1. present in ≥ 10% of reference genomes; 2. average copy number ≤ 1.2; and 3. average predicted protein length > 100 amino acid residues. For each core gene, matching proteins in viralRefSeq references and vOTUs (this study) were identified and predicted protein amino acid sequences aligned with Clustal Omega v1.2.4[10]. Sequences with predicted genome completeness ≤ 85% (estimated via CheckV) were omitted to minimise false weighting from missing genes due to genome sequence incompleteness. Aligned proteins were trimmed to remove columns (amino acid positions) represented in <50% of genomes prior to concatenation (with gaps introduced where proteins were missing from a genome). Finally, genomes with amino acid representation of <5% total concatenated alignment length were removed, resulting in a final concatenated protein alignment matrix of 13,415 columns (amino acid positions) x 4,481 genomes, with 85% missing data (missing data value comparable to that of Low and colleagues (84%)[6]). Phylogenetic trees based on the concatenated alignment were generated via IQ-TREE v2.2.2.2[11] using ModelFinder (with VT+F+I+G4 selected)[12], and with ultrafast bootstrap (nmax = 1000 replicates)[13]. Trees were visualised using iTOL[14].

An alternative set of putative core genes were also identified for all high quality *Caudoviricetes* sequences (based on the provided taxonomy) in the IMG/VR database (**Table S6**) following the method outlined above with the following modifications: IMG/VR sequences were partitioned 100 ways (partition.sh from BBMap suite of tools v38.95[15]), and genes predicted via prodigal-gv v2.9.0[16,17]. For each partition, hmmsearch against HMM profiles from the VOGdb database was performed with -Z 24911692 (the count of all predicted proteins from the IMG/VR dataset across all partitions) and -E 1e-3. Separate concatenated protein alignments of core genes and phylogenetic trees were generated based on each of the RefSeq and IMG/VR core genes sets, as per the method outlined above.

*Identification of putative prokaryote hosts*

Putative prokaryote hosts for Waiwera estuary vOTUs were predicted using a combination of approaches, including CRISPR-spacer, tRNA, and genome homology via pairwise blast searches (BLAST v2.13.0)[18], analysis of vOTUs co-binned with prokaryote MAGs, oligonucleotide frequency similarity via VirHostMatcher v1.0.0[19], and machine learning-based methods RaFAH v0.3[20] and HostG (accessed 06 Dec 2021)[21]. These six methods were applied to the non-dereplicated MAG dataset and to all viral contigs, which were later summarised by vOTU cluster. CRISPR-spacers were extracted from filtered and trimmed metagenomics sequencing reads using crass v1.0.1[22] with the default parameters. Spacer sequences were searched against all non-dereplicated prokaryote MAGs (pre-dRep MAGs; n = 905) and viral contigs using blastn with the parameters -dust no -word_size 7. Viral contigs and MAGs that shared perfect matches to two or more unique spacer sequences (zero mismatches or gaps) were considered as a putative virus-host match. To provide more confidence in virus-host predictions based on CRISPR matching, CRISPR regions were also searched within all MAGs via CRISPRDetect v2.4[23] with -array_quality_score_cutoff 1. tRNA sequences were identified in prokaryote MAGs and vOTUs from the Waiwera estuary with Aragorn v1.2.38[24], followed by pairwise blastn with the parameters -num_alignments 5 -dust no, and filtering to retain the top three matches with ≥ 90% nucleotide identity over ≥ 90% sequence length. Genome homology between MAGs and vOTUs was compared via blastn with filtering to retain the top three matches with nucleotide identity ≥ 70%, bit score ≥ 50%, and evalue ≤ 0.001. Contigs predicted to contain viral sequence were searched within prokaryote MAGs to identify any viruses that were co-binned with individual MAGs. Co-binned vOTUs also predicted to be prophage (based on trimming conducted by VIBRANT, VirSorter2, or CheckV) were identified as candidate integrated prophage within the host MAG. For VirHostMatcher, the Waiwera Estuary prokaryote MAG dataset (this study) was provided as the database of potential hosts, with filtering to retain the top five hits with dissimilarity score ≤ 0.25. RaFAH and HostG were run using the default settings, with additional filtering for HostG to retain predictions with softmax scores ≥ 0.8.
